# The Fanconi anemia core complex promotes CtIP-dependent end-resection to drive homologous recombination at DNA double-strand breaks

**DOI:** 10.1101/2023.09.05.556391

**Authors:** Bert van de Kooij, Fenna J. van der Wal, Magdalena B. Rother, Pau Creixell, Merula Stout, Wouter Wiegant, Brian A. Joughin, Julia Vornberger, Marcel A.T.M. van Vugt, Matthias Altmeyer, Michael B. Yaffe, Haico van Attikum

## Abstract

Homologous Recombination (HR) is a high-fidelity repair mechanism of DNA Double-Strand Breaks (DSBs), which are induced by irradiation, genotoxic chemicals or physiological DNA damaging processes. DSBs are also generated as intermediates during the repair of interstrand crosslinks (ICLs). In this context, the Fanconi anemia (FA) core complex, which is effectively recruited to ICLs, promotes HR-mediated DSB-repair. However, whether the FA core complex also promotes HR at ICL-independent DSBs remains controversial. Here, we identified the FA core complex members FANCL and Ube2T as HR-promoting factors in a CRISPR/Cas9-based screen with cells carrying the DSB-repair reporter DSB-Spectrum. Using isogenic cell-line models, we validated the HR-function of FANCL and Ube2T, and demonstrated a similar function for their ubiquitination-substrate FANCD2. We further show that FANCL and Ube2T are directly recruited to DSBs and are required for the accumulation of FANCD2 at these break sites. Mechanistically, we demonstrate that FANCL ubiquitin ligase activity is required for the accumulation of the nuclease CtIP at DSBs, and consequently for optimal end-resection and Rad51 loading. CtIP overexpression rescues HR in FANCL-deficient cells, validating that FANCL primarily regulates HR by promoting CtIP recruitment. Together, these data demonstrate that the FA core complex and FANCD2 have a dual genome maintenance function by promoting repair of DSBs as well as the repair of ICLs.

## Introduction

DNA double-strand breaks (DSBs) are dangerous DNA lesions that separate a chromosome into two fragments. If left unrepaired, DSBs can result in mitotic mis-segregation of the broken chromosome and subsequent aneuploidy. Hence, efficient repair of DSBs is essential to maintain genome stability. This is ensured by the collective activity of a variety of DSB-repair pathways, including homologous recombination (HR)^1^. DSB-repair by HR is initiated by end-resection, which involves nuclease-mediated strand-removal at the DSB-ends to generate 3’ single-strand overhangs^2^. These overhangs are bound by the recombinase protein Rad51 that drives invasion of the 3’ DNA overhang into a homologous DNA region, most often the sister chromatid ^3^. Subsequently, multiple HR sub-pathways can be distinguished, all of which involve extension of the DNA overhang on the homologous DNA, followed by untangling of the recombination intermediate and completion of repair.

HR-initiation by end-resection is a tightly coordinated process^2^, which starts with binding of the Mre11-Rad50-Nbs1 (MRN) complex to DSB-ends ^2^. Mre11 has endonuclease activity, which is used to nick the DNA adjacent to the DSB, as well as exonuclease activity, which is used to resect the DNA from the nick towards the DSB-end^2^. The endonuclease activity of Mre11 is strongly promoted by its co-factor CtIP^4,5^. CtIP is a central regulator of end-resection that, besides promoting Mre11, can also enhance the activity of the nuclease DNA2, which acts downstream of Mre11 to processively resect the DNA in the 5’ to 3’ direction^6^. CDK-mediated phosphorylation of CtIP restricts its activity to the S/G2 cell-cycle phases, thus synchronizing end-resection with the presence of the sister chromatid^7,8^. CtIP interacts with the Mre11-Rad50-Nbs1 (MRN) complex, as well as with the HR-factor BRCA1^5,9–11^. However, neither of these interactions are essential for the recruitment of CtIP to DSBs, indicating that this can be mediated by additional signals at the break-site^12–14^.

In addition to HR, DSBs can be repaired by canonical Non-Homologous End-Joining (c-NHEJ), alternative End-Joining (a-EJ), or Single-Strand Annealing (SSA)^1^. Whereas c-NHEJ involves very minimal end-processing prior to ligation, a-EJ and SSA require end-resection to reveal regions of homology that can base-pair to join the two opposing DSB-ends. Repair by c-NHEJ, a-EJ and SSA effectively reconnects the broken chromosome fragments, yet mostly at the expense of mutations at the break junction. In contrast, HR is generally considered to faithfully restore the original DNA sequence. Hence, HR forms a barrier against mutagenesis and chromosomal alterations, and as such is an important tumor suppressor pathway. In agreement, a high frequency of tumors, in particular those derived from breast and ovarian tissue, are HR-deficient due to germline or somatic mutations in HR-genes like *BRCA1* or *BRCA2*^15,16^.

Considering the important genome maintenance function of HR, we sought to identify novel genes involved in DSB-repair by this pathway. To this end, we performed a targeted CRISPR-based genetic screen in a cell-line carrying the DSB-repair reporter DSB-Spectrum^17^. As we show here, the results from this screen suggested that the E2 ubiquitin conjugase Ube2T and the E3 ubiquitin ligase FANCL function as HR-promoting factors. Ube2T and FANCL are both part of the multi-member Fanconi anemia (FA) core complex that plays a well-characterized role during the repair of DNA interstrand crosslinks (ICLs)^18^. Inactivating mutations in the genes encoding the FA core complex members, or downstream ICL-repair factors, are all associated with the hereditary disorder Fanconi anemia^19^. The FA core complex recognizes crosslinked DNA and subsequently ubiquitinates the FANCD2/FANCI heterodimer^18^. This is an essential step to recruit nucleases that remove the ICL, but also to promote the downstream repair steps, including HR-mediated repair of the DSB that is generated following ICL removal.

The FA core complex and FANCD2/FANCI have been suggested to also promote HR at DSBs that are generated independently of ICL-repair^20–23^. However, the relevance of this function has been questioned, as the HR-phenotypes observed upon depletion of FA factors were mild or even absent in some studies^22,24–28^. Moreover, mechanistic insight into the DSB-repair function of the FA core complex is lacking. Here, we validate the HR-promoting function of Ube2T and FANCL observed in our CRISPR-based genetic screen using orthogonal approaches in a variety of knock-out cell-lines and isogenic control cell-lines. Moreover, we show that Ube2T and FANCL activity are required for optimal CtIP-dependent end-resection at DSBs, providing a mechanistic explanation for their HR-promoting function. Together, our data indicate that the FA core complex and FANCD2 not only promote ICL-repair, but are also *bona fide* DSB-repair factors that act during an initial and essential stage of HR.

## Results

### A DSB-Spectrum reporter screen identifies members of the FA core complex as HR-promoting factors

To identify novel genes that drive error-free DSB-repair, a genetic screen was performed using DSB-Spectrum_V2, a genomic DSB-repair reporter that we had previously created to distinguish mutagenic repair from HR (Fig. 1A)^17^. The reporter consists of a functional Blue Fluorescent Protein (BFP) gene separated by an ∼3 kilobase region from a promotorless and truncated Enhanced Green Fluorescent Protein gene (EGFP, hereafter referred to as GFP). In this system, a single DSB is generated by targeting Cas9 to the BFP gene at a site adjacent to the chromophore-determining amino acids. Mutagenic repair of the DSB, for example by c-NHEJ, will disrupt the BFP gene, resulting in loss of fluorescence (Fig. 1A). Alternatively, repair of the DSB by HR using the downstream, highly homologous truncated GFP gene as a repair template will replace the BFP serine-66 and histidine-67 encoding triplets with threonine and tyrosine-encoding triplets of the truncated GFP gene, causing BFP-to-GFP conversion^29^. Thus, mutagenic repair or HR-mediated repair of a DSB is detected by total loss of fluorescence or by conversion from BFP to GFP expression, respectively.

**Figure 1.**
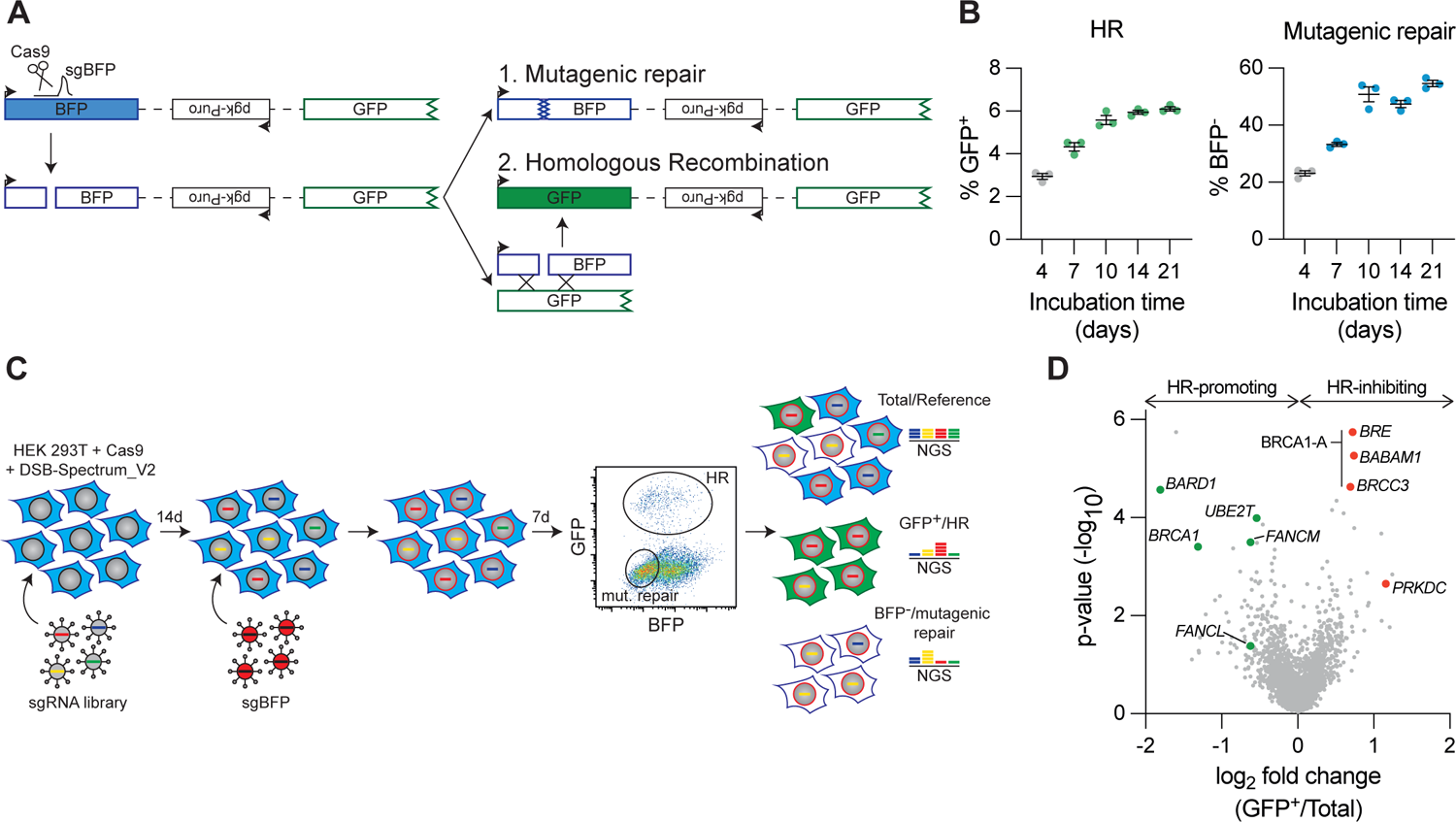
A targeted CRISPR screen in DSB-Spectrum reporter cells identifies FANCM, Ube2T and FANCL as HR-promoting factors. **(A)** Schematic of the DSB-Spectrum_V2 reporter. Adpated from van de Kooij *et al.*, 2022. **(B)** HEK 293T+Cas9+DSB-Spectrum_V2 cells were lentivirally infected to express mCherry and the BFPsg targeting the DSB-Spectrum_V2 reporter locus. Next, at indicated time points, BFP and GFP expression was analyzed by flow cytometry. Depicted is the mean±SEM of a technical triplicate. **(C)** Schematic displaying the CRISPR screen lay-out in HEK 293T+Cas9+DSB-Spectrum_V2 cells. **(D)** Volcano plot showing the gene targets of sgRNAs that were either enriched or depleted from the GFP^+^ HR population as compared to the reference population. *BRCA1/BARD1* and *Ube2T/FANCM/FANCL* are indicated in green as HR-promoting factors. *BRE/BABAM1/BRCC3*, all members of the BRCA1-A complex, and *PRKDC* are indicated in red as HR-inhibiting factors.

To generate a reporter cell-line suitable for genetic screening, HEK 293T cells stably expressing Cas9 were lentivirally transduced to introduce a copy of DSB-Spectrum_V2 into the genome. A single-cell clone was expanded to create a homogeneous HEK 293T DSB-Spectrum_V2 cell-line. To validate this cell-line and establish the kinetics of repair, a BFP-targeting sgRNA or AAVS1-targeting control sgRNA was introduced by lentiviral transduction, and BFP and GFP expression were analyzed by flow cytometry at various time points after transduction. Cells expressing the control sgRNA remained BFP-positive and GFP-negative throughout the experiment (Fig. S1A). In contrast, in cells expressing the BFP-targeting sgRNA, both BFP-negative/GFP-negative and BFP-negative/GFP-positive populations could be detected at four days after transduction, indicating mutagenic and HR-mediated DSB-repair, respectively (Fig. 1B and S1A). The mutagenic and HR repair populations gradually increased in size until reaching a plateau at 10 days after transduction (Fig. 1B and S1A). As we had previously validated that these distinct fluorescent populations are the consequence of mutagenic DSB-repair and HR, respectively^17^, we concluded that the newly-generated DSB-Spectrum_V2 reporter cell-line is functional.

To identify genes that modulate different DNA repair phenotypes, a CRISPR-based genetic screen was performed (Fig. 1C). A custom-generated single-guide RNA (sgRNA) library targeting 2,760 genes, with four sgRNAs per gene, was introduced into the DSB-Spectrum_V2 reporter cells by lentiviral transduction (Table S1). The targeted genes encoded kinases and phosphatases, ubiquitin and SUMO modifiers, and factors that read, write or remodel chromatin. Cells were cultured for 14 days to allow editing of the target genes and depletion of their protein products, after which the BFP-targeting sgRNA was introduced by lentiviral infection to generate a DSB within the DSB-Spectrum_V2 reporter. At seven days after BFP-targeting, the BFP-negative/GFP-positive HR and BFP-negative/GFP-negative mutagenic repair populations were harvested by Fluorescence-Activated Cell Sorting (FACS). In addition a sample from the total population was collected for use as a reference. sgRNA counts in all samples were determined by Illumina-sequencing.

As an initial quality control measure, we determined the level of depletion of sgRNAs targeting essential genes^30^. The majority of sgRNAs targeting essential genes were strongly depleted from the total surviving population relative to the input library, validating that CRISPR-based gene editing was sufficient to permit a measurable phenotype in our system (Fig. S1B). As a second quality control measure, we assessed how sgRNAs targeting known DSB-repair factors behaved. A clear depletion of sgRNAs targeting the HR-promoting genes *BRCA1* and *BARD1* was observed in the GFP-positive HR population compared to the reference population (Fig. 1D, table S2). In contrast, sgRNAs targeting *PRKDC* or the members of the BRCA1-A complex, which both inhibit HR, were strongly enriched in the HR population (Fig. 1D, table S2). Thus, this screening approach can faithfully detect known factors that promote or impair HR. Theoretically, analysis of sgRNA levels in the GFP-negative/BFP-negative population should allow for the identification of factors involved in mutagenic DSB-repair. However, we were not able to validate this branch of the screen, as no enrichment or depletion of known DSB-repair factors was observed in this mutagenic repair population, probably as a consequence of re-balancing repair by multiple mutagenic pathways (table S2)^17^. Considering this result, and based on our primary interest in HR, we therefore decided to focus on the HR-branch of this screen.

The custom-generated library used for our screen contained sgRNAs targeting three FA genes: *Ube2T*, *FANCL*, and *FANCM*. Interestingly, for all three FA genes we observed strong depletion of their cognate sgRNAs from the HR-population, with very low p-values for *Ube2T*- and *FANCM-*targeting sgRNAs (Fig. 1D, table S2). To determine the generalizability of these hits, we compared our results with those of a recently published CRISPRi screen in K562 cells aimed at identifying regulators of HR using an ectopically provided dsDNA donor^23^. Although we observed remarkably limited correlation between the two datasets, *Ube2T*, *FANCL* and *FANCM* were among the few candidates identified as HR-promoting factors in both screens (Fig S1C). Together, these data therefore strongly suggest a DSB-repair function for the FA core complex that is independent of its role in ICL-repair.

### Optimal HR requires the expression and ubiquitin ligase activity of FANCL and Ube2T

FANCM has been suggested to promote short-tract gene conversion during HR by dissolving recombination intermediates like double Holliday junctions^31–33^. This HR-function of FANCM, which is independent of the FA core complex, could explain why FANCM was identified as an HR-promoting factor in our screen. In contrast, the functions of Ube2T and FANCL in HR are largely underexplored. We therefore decided to focus on these two FA proteins in more detail. The results from the CRISPR screen were first validated by targeted depletion of Ube2T or FANCL in HR-reporter assays using HEK 293T cells carrying the reporter DSB-Spectrum_V3, a next-generation variant of DSB-Spectrum_V2. This V3 reporter contains an mCherry gene between the BFP and truncated GFP genes to allow mutagenic DSB-repair resulting from direct end-joining to be distinguished from that resulting from SSA (Fig. S2A)^17^. DSB-repair by SSA results in deletion of the mCherry gene, which can be monitored by flow cytometry, in conjunction with repair by HR (GFP-positive) and mutagenic end-joining (mut-EJ; BFP-negative/ mCherry-positive population of cells). Monoclonal *Ube2T* knock-out (*Ube2T*^KO^) cell-lines were generated using CRISPR-technology. As seen in figure 2A, two Ube2T antibody-responsive bands were detected on Western blots from the Ube2T wild-type (WT) parental control cell-line (Con.). The lower band could still be detected in *Ube2T*^KO^ clone 3.3, albeit with lower intensity, suggesting that this may be a partial knock-out (Fig. 2A, see Fig. S4 for uncropped blots). In clone 4.1 neither of the two Ube2T bands was detected, indicating that this clone is a complete Ube2T knock-out.

**Figure 2.**
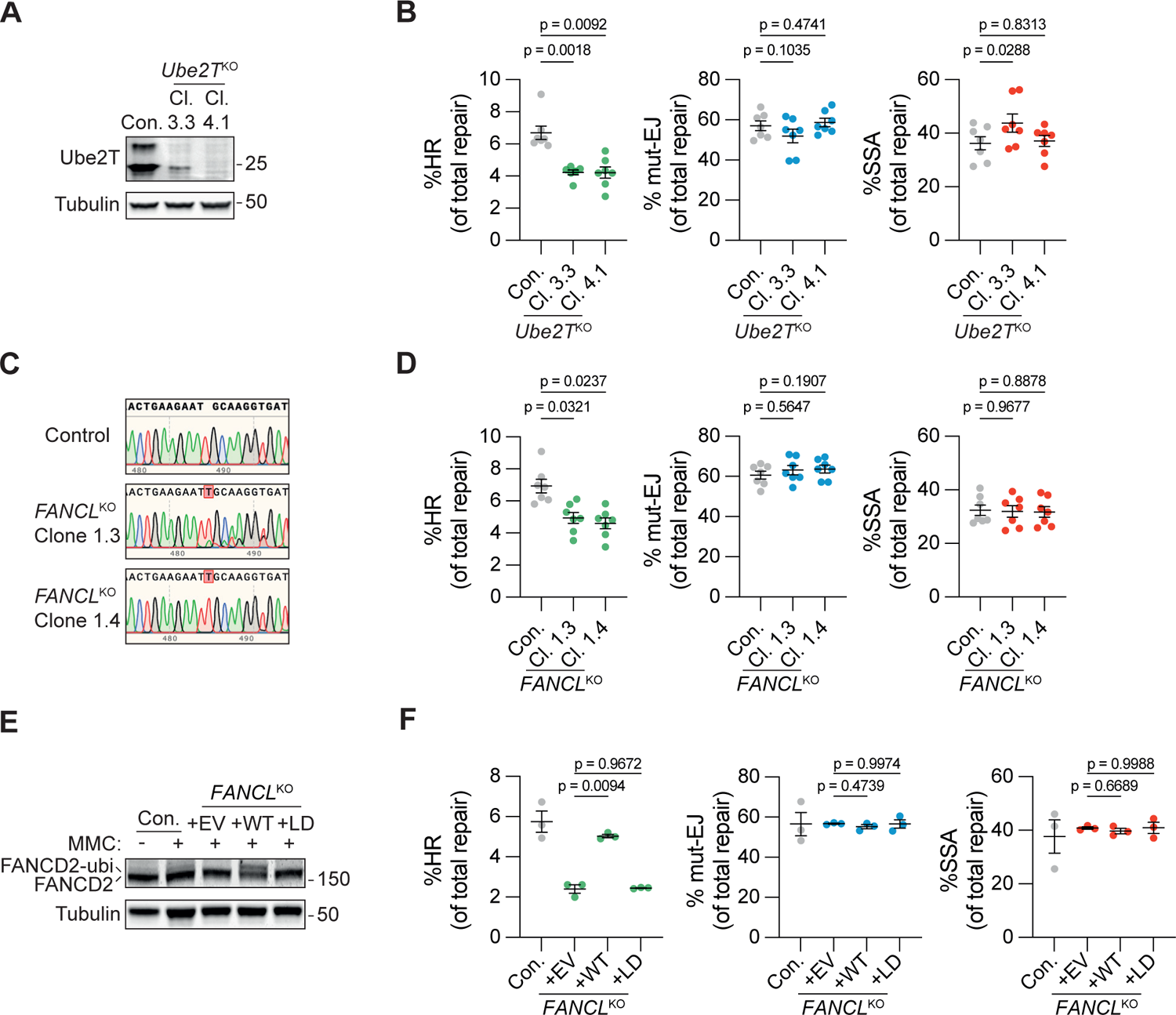
Ube2T and FANCL promote HR-mediated repair of Cas9-induced DSBs. **(A)** Ube2T expression was analyzed by western blot in HEK 293T+DSB-Spectrum_V3 *Ube2T*^KO^ clones 3.3 and 4.1, and in the parental control. **(B)** Indicated HEK 293T+DSB-Spectrum_V3 cell-lines were transfected to express Cas9 and either a control sgRNA, or the sgRNA targeting the BFP gene in the reporter locus. Next, cells were analyzed by flow cytometry to determine the frequency of repair by each of the three indicated pathways (n=7; mean±SEM; One-way ANOVA+Dunnett’s multiple comparison). **(C)** DNA sequence alignment of the *FANCL* sgRNA target site in unedited parental control cells and the HEK 293T+DSB-Spectrum_V3 *FANCL*^KO^ clones. Depicted are representative sequence chromatograms, red shaded boxes indicate deviations in the DNA sequence of the *FANCL*^KO^ clone compared to control. **(D)** As in panel B, now analyzing *FANCL*^KO^ cells. Note that for four biological repeats the same control was shared between panel B and this panel (n=7; mean±SEM; One-way ANOVA+Dunnett’s multiple comparison). **(E)** HEK 293T+DSB-Spectrum_V3 *FANCL*^KO^ clones were transduced with an empty vector (EV), FANCL wild-type cDNA (WT), or FANCL Ligase-Dead cDNA (LD), and treated with Mitomycin C (MMC, 500μM) for 24h. Next, FANCD2 ubiquitination was analyzed by western blot. **(F)** As in panel B, now analyzing the HEK 293T+DSB-Spectrum_V3 *FANCL*^KO^ cells described in panel E (n=3; mean±SEM; One-way ANOVA+Dunnett’s multiple comparison).

The *Ube2T*^KO^ and control cells were transfected to express Cas9, the BFP sgRNA, and the iRFP(670) protein as a marker for transfected cells, and the frequency of HR, mut-EJ and SSA was determined by flow cytometry 72-96h later. DSB-repair by HR was significantly reduced in both *Ube2T*^KO^ clones as compared to the control cells (Fig. 2B). In contrast, the level of mut-EJ was similar between the control and *Ube2T*^KO^ cell-lines. SSA was mildly, but significantly, increased in *Ube2T*^KO^ clone 3.3, as compared to the control cells. Of note, this SSA phenotype was not shared by either *Ube2T*^KO^ clone 4.1, or the *FANCL*^KO^ clones described below. We therefore conclude that Ube2T specifically promotes DSB-repair by HR, thus validating the results from the screen.

HEK 293T DSB-Spectrum_V3 cell-lines in which FANCL was genetically deleted were generated next. The levels of FANCL protein could not be determined due to the absence of any functional anti-FANCL antibody. Therefore, the knock-out status of the selected *FANCL*^KO^ clones was confirmed by sequence analysis of the genomic target site, which identified out-of-frame mutations in all alleles (Fig. 2C). HR was significantly reduced in these *FANCL*^KO^ cell-lines compared to control, as shown by DSB-Spectrum_V3 reporter assays, whereas the levels of mut-EJ and SSA were unaffected (Fig. 2D). To confirm that the observed phenotype was the specific result of loss of FANCL function, wild-type FANCL cDNA (+WT), or an empty vector control (+EV), was re-introduced into *FANCL*^KO^ clone 1.4. In addition, we introduced a FANCL mutant carrying a C307A mutation that abrogates its ligase activity (Ligase-Dead, +LD)^34^. To first validate rescue of FANCL function, cell-lines were treated with the DNA cross-linking agent mitomycin C (MMC) and the ubiquitination of the FANCL substrate FANCD2 was monitored by Western blotting. FANCD2 ubiquitination, indicated by the appearance of a slower migrating FANCD2 species, was clearly observed in the *FANCL*^KO^ cells reconstituted with FANCL WT, but not in the cells reconstituted with either the EV or the FANCL LD mutant (Fig. 2E), indicating that only the +WT cells express functional ligase-competent FANCL. The HR phenotype in the different FANCL cell-lines was examined next using the DSB-Spectrum_V3 reporter assay. In agreement with the data shown above, the *FANCL*^KO^ cells (+EV) showed strongly impaired HR compared to the control cells, which was rescued by re-expression of FANCL WT, but not by re-expression of the FANCL LD mutant (Fig. 2F). Taken together, these data indicate that UbeT2 and FANCL expression, as well as FANCL ligase activity, are required for efficient HR.

### Loss of Ube2T, FANCL or FANCD2 sensitizes cells to PARP inhibitor treatment

To further validate the HR-function of Ube2T and FANCL, we utilized an alternative assay orthogonal to the DSB-Spectrum_V3 reporter assays, and explored cells other than HEK 293T^34,35^. Clonally derived Ube2T and FANCL knock-out cell-lines were generated in a U-2 OS background, and validated by Western blotting and sequence analysis for two *Ube2T*^KO^ and two *FANCL*^KO^ clones, respectively (Fig. 3A, S2B). Of note, a single Ube2T species was detected by Western blot analysis of the U-2 OS lysates, in contrast to the doublet that we consistently observed in lysates from HEK 293T cells. The knock-out status of the *Ube2T*^KO^ and *FANCL*^KO^ cell-lines was further validated by the absence of MMC-induced FANCD2 ubiquitination in these cell-lines (Fig. 3A, B).

**Figure 3.**
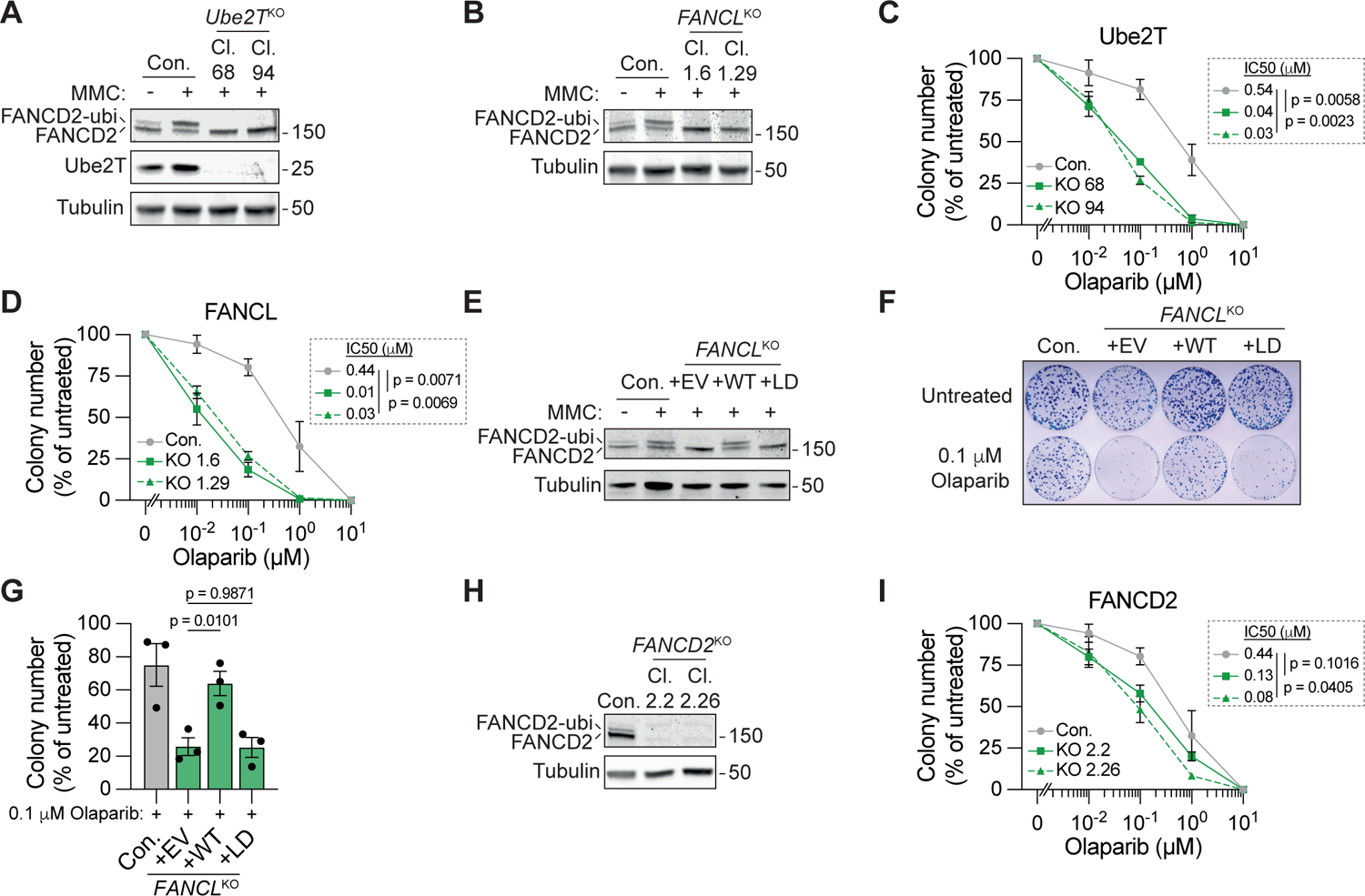
Loss of Ube2T, FANCL and FANCD2 sensitizes to PARP inhibitor-induced toxicity. **(A)** Indicated U-2 OS *Ube2T*^KO^ clones, as well as the parental control, were treated with Mitomycin C (MMC, 500 μM) for 24h. Next, FANCD2 ubiquitination status and protein levels of Ube2T were analyzed by western blot. **(B)** As in panel A, but now for U-2 OS *FAN-CL*^KO^ clones. **(C and D)** U-2 OS cells, either wild-type control (Con.) or knock-out for Ube2T (panel C) or FANCL (panel D), were treated with olaparib for 14 days. Cell viability was assed by clonogenic survival. Note that the 0 μM value was added manually to the X-axis. Inset shows mean IC50 and p-value (n=3, expect for the 0.1 μM and 10 μM concentration in panel C for which n=2; mean±SEM; Ratio paired t-test). **(E)** As in panel B, now for *FANCL*^KO^ cells that were transduced with an empty vector (EV), FANCL wild-type cDNA (WT), or FANCL ligase-dead cDNA (LD). **(F and G)** Indicated cell-lines were treated with 0.1 μM olaparib, or left untreated, for 14 days. Cell viability was assessed by clonogenic survival. Panel F shows a representative picture of Methylene Blue-stained colonies, panel G shows the quantification (n=3; mean±SEM; One-way ANOVA+Dunnett’s multiple comparison). **(H)** As in panel B, but now for untreated *FANCD2*^KO^ clones. **(I)** As in panels C and D, but for FANCD2^KO^ cells (n=3; mean±SEM; Ratio paired t-test).

PARP inhibitor (PARPi) treatment is commonly used as a proxy for HR-pathway activity because HR-defects strongly sensitize cells to the toxic effects of PARP inhibition^35,36^. The *Ube2T*^KO^ and *FANCL*^KO^ clones, as well as control cells, were therefore treated with varying doses of the PARPi olaparib, and clonogenic outgrowth was measured to asses HR activity. All *Ube2T*^KO^ and *FANCL*^KO^ cell-lines were significantly more sensitive to PARPi-induced viability loss than the control cell-line (Fig. 3C, D). To further assess the role of FANCL ligase activity in the olaparib response, *FANCL*^KO^ clone 1.6 was reconstituted with FANCL WT, the LD mutant, or EV control. As expected, MMC-induced FANCD2 ubiquitination was observed in the FANCL WT, but not the EV or FANCL LD cell-lines, validating that re-expressed FANCL WT was functional (Fig. 3E). Moreover, re-expression of FANCL WT, but not the FANCL LD mutant, rescued the increased PARPi sensitivity of the *FANCL*^KO^ clone (Fig. 3F, G). Hence, loss of Ube2T or FANCL expression or activity sensitizes cells to PARPi treatment, consistent with an HR-promoting function for both of these factors.

During repair of interstrand crosslinks, FANCD2 is the downstream substrate of Ube2T and FANCL. To examine whether FANCD2 functions in DSB-repair by HR as well, we generated *FANCD2*^KO^ cell-lines in the U-2 OS background and measured PARPi sensitivity (Fig. 3H, I). As seen with Ube2T and FANCL, loss of FANCD2 similarly reduced clonogenic outgrowth after olaparib treatment (Fig. 3I). Notably, loss of FANCD2, or of FANCL or Ube2T, did not significantly affect cell-cycle distribution (Fig. S2C). Thus, the PARPi sensitivity phenotype of cells following loss of FANCD2, FANCL or Ube2T is unlikely to be caused by changes in cell cycle distribution. Instead, these results suggest that FANCD2, Ube2T, and FANCL cooperate to directly promote HR-repair of DSBs.

### The FA core complex is directly recruited to DSBs, where it promotes FANCD2 accumulation

To determine whether Ube2T and FANCL directly promote HR at DSB-sites, we examined their recruitment to sites of DNA damage using laser micro-irradiation. U-2 OS cell-lines stably expressing GFP-tagged FANCL or Ube2T, as well as a control cell-line expressing GFP with a nuclear localization signal (NLS) were generated, pre-treated with BrdU and exposed to UV-A laser micro-irradiation to generate localized stripes of DNA damage, predominantly DSBs^37^. Subsequently, cells were fixed and analyzed by immunofluorescence microscopy to study the enrichment of Ube2T or FANCL at the sites of DNA damage. A staining to detect phosphorylated histone H2AX (γH2AX) was taken along to identify the DNA damage stripes. Whereas GFP-NLS was equally distributed throughout the nucleus, both FANCL and Ube2T were significantly enriched in the DNA damage stripes (Fig. 4A, B). Notably, enrichment was stronger for FANCL than for Ube2T, consistent with a scaffolding function for FANCL and an enzymatic function for Ube2T (Fig. 4A, B). Hence, both FANCL and Ube2T are recruited to UV-A laser-induced DSBs.

**Figure 4.**
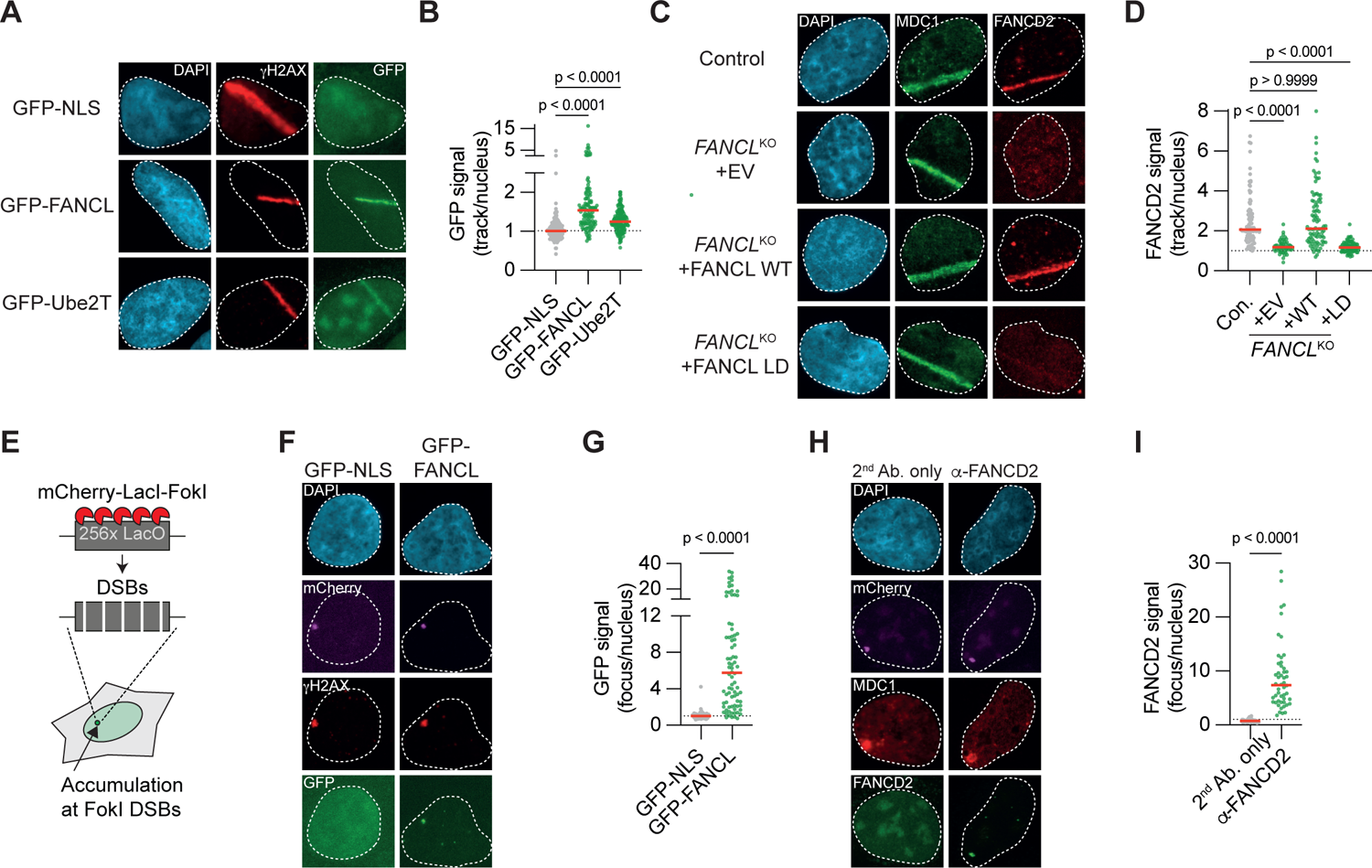
FANCL, Ube2T and FANCD2 are recruited to DNA Double-Strand Breaks. **(A, B)** U-2 OS cells expressing either GFP-NLS as a control, or GFP-tagged FANCL or Ube2T were exposed to UV-A laser micro-irradiation. Next, cells were fixed and analyzed by immunofluorescence microscopy. Laser-induced DNA damage tracks were identified by α-γH2AX staining. Panel A shows representative images, panel B shows the quantification of a representative experiment. Dotted line is set at 1 (*i.e.* no recruitment to the track), red lines indicate median (n=2; one-way ANOVA with post-hoc Kruskal-Wallis). **(C, D)**. As in panels A, B, but now analyzing recruitment of endogenous FANCD2 to UV-A laser induced DNA damage tracks in U-2 OS *FANCL*^KO^ cell-lines (EV=empty vector). DNA damage tracks were identified by α-MDC1 staining. Panel C shows representative images, panel D shows the quantification of a representative experiment. Red lines indicate the median (n=2; one-way ANOVA with post-hoc Kruskal-Wallis). **(E)** Cartoon schematic of a DSB recruiment assay in U2OS 2-6-3 cells. In short, LacI-fused mCherry-tagged FokI is tethered to an array of LacO repeats where it generates a large number of DSBs. DSB-repair factors accumulate at these DSBs and form microscopically discernable foci. Adapted from Singh *et al*., 2021 **(F, G)** Accumulation of GFP-NLS or GFP-FANCL at γH2AX-marked FokI-generated DSBs in U-2 OS 2-6-3 cells was assessed by immunofluorescence microscopy. Panel F shows representative images, panel G shows the quantification of a representative experiment. Red lines indicate the median (n=2; Mann-Whitney test). **(H,I)** As in panels F and G but now analyzing endogenous FANCD2 recruitment to MDC1-marked FokI-induced DSBs (n=2; Mann-Whitney test).

FANCD2 has previously been shown to accumulate at DNA damage sites upon DSB inducing treatments^38,39^. To examine whether FANCD2 recruitment to DSBs was dependent on FANCL, we assessed FANCD2 localization using laser micro-irradiation assays in the reconstituted U-2 OS *FANCL*^KO^ cells described above. Strong enrichment of endogenous FANCD2 was observed in control cells at UV-A laser-induced DNA damage stripes marked by the γH2AX-binding protein MDC1 (Fig. 4C). Importantly, no FANCD2 enrichment was detected in the *FANCL*^KO^ cells reconstituted with empty vector (Fig. 4C, D). Re-expression of FANCL WT, but not of the FANCL LD mutant, however, rescued the FANCD2 recruitment to sites of DNA damage. Hence, FANCL expression and ligase activity are required for the accumulation of FANCD2 at UV-A laser induced DSBs. To extend these finding to other members of the FA core complex, we studied the patient-derived head and neck squamous cell carcinoma cell-line VU1131, which is deficient for FANCC^40^. No recruitment of FANCD2 to DNA damage stripes was observed in these cells. However, re-expression of FANCC completely rescued the FANCD2 recruitment defect (Fig. S2D, E). These data indicate that Ube2T and FANCL act as part of the FA core complex to promote FANCD2 recruitment to DSBs.

UV-A laser micro-irradiation in BrdU-treated cells predominantly causes DSBs, but a minority of other DNA lesions can also be generated that could be at least partially responsible for the accumulation of FA factors at the irradiated sites. To exclude this possibility, the recruitment of FA factors to nuclease-induced DSBs was examined using an alternative approach. GFP-FANCL and GFP-Ube2T expressing cell-lines were generated in U-2 OS 2-6-3 cells that contain an mCherry-tagged, LacI-fused FokI nuclease and a genomically integrated LacO array^41^. Upon treatment with tamoxifen and Shield-1, the FokI nuclease is stably expressed, translocates to the LacO array, and generates a multitude of DSBs within the LacO array, resulting in the local accumulation of DSB-repair factors (Fig. 4E)^41^. Following treatment of our cell-lines with tamoxifen and Shield-1, a single distinct γH2AX focus was observed that co-localized with the mCherry-FokI nuclease (Fig. 4F). In most cells, the GFP-FANCL signal was strongly enriched in these FokI foci (Fig. 4F, G). GFP-Ube2T was also significantly enriched in FokI foci, in contrast to GFP-NLS, which served as a negative control (Fig. S2F, G). Notably, similar to what was observed at UV-A laser-induced DNA damage sites, accumulation of Ube2T at FokI-induced DSB-sites was substantially less than the accumulation of FANCL (Fig. 4B, G, S2G), in agreement with their proposed enzymatic and structural roles, respectively. Furthermore, strong accumulation of endogenous FANCD2 at FokI-induced DSBs was observed (Fig. 4H, I). Collectively, these results demonstrate that Ube2T, FANCL and FANCD2 are recruited to *bona fide* DSBs, suggesting that they act directly at DNA break sites to promote repair by HR.

### FANCL promotes DSB end-resection

We next explored the mechanism by which FANCL and Ube2T promote DSB-repair by HR. Since FANCD2 has previously been suggested to function in DNA end-resection ^42^, we hypothesized that this may require the upstream activity of FANCL/Ube2T. To test this, U-2 OS *FANCL*^KO^ cells were exposed to ionizing radiation (IR), and the accumulation of serine 4/8 phosphorylated RPA (pRPA) at DNA damage-induced foci was measured as a proxy for the presence of resected ssDNA^43^. In addition, a short pulse of EdU was administered prior to the IR-exposure to allow for the detection of S-phase cells, which are HR-prone. The formation of IR-induced pRPA foci was strongly reduced in the *FANCL*^KO^ cells as compared to the control cells (Fig. 5A, B, S3A). A similarly strong reduction in pRPA foci number was observed in *Ube2T*^KO^ and *FANCD2*^KO^ cells (Fig. S3B-D). The reduction in nuclear pRPA levels in the *FANCL*^KO^ cells could be completely rescued by re-expression of FANCL WT (Fig. 5C, S3E). Re-expression of the FANCL LD mutant also increased nuclear pRPA levels, albeit to a level that was still significantly lower than in the control cells (Fig. 5C).

**Figure 5.**
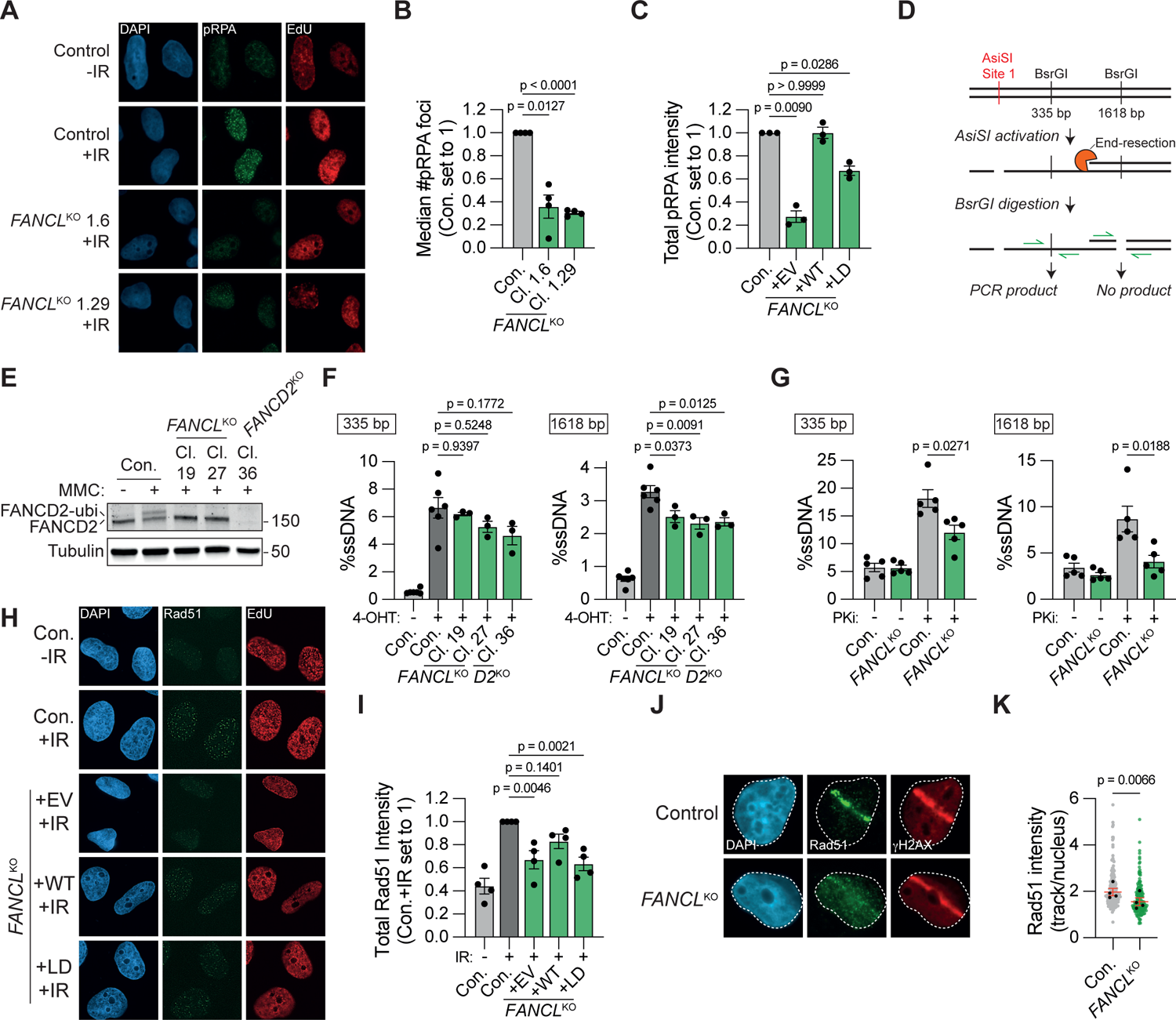
FANCL promotes end-resection at DNA double-strand breaks. **(A, B)** U-2 OS *FANCL*^KO^ and wild-type control cells were exposed to 10 Gy ionizing radiation (IR), followed by IF microscopy to detect foci containing S4/8 phosphorylated RPA (pRPA) in S-phase (EdU+) nuclei. Panel A shows representative images, panel B shows the quantification (n=4; mean±SEM; one-way ANOVA with post-hoc Dunnett’s). **(C)** As in panel B, but now plotting total pRPA intensity per nucleus in *FANCL*^KO^ cells reconstituted with FANCL WT or LD (n=3; mean±SEM; one-way ANOVA with post-hoc Dunnett’s). **(D)** Schematic of the qPCR-based quantification of end-resection in AsiSI cells. **(E)** Western blot of MMC-treated (500 μM, 24h) U-2 OS AsiSI cells. **(F)** Quantification by qPCR of single-strand DNA (ssDNA) at 335 bp or 1618 bp distance from a defined AsiSI-induced DSB (n≥3; one-way ANOVA with post-hoc Dunnett’s). **(G)** As in panel F, but now including treatment with the DNA-PKcs inhibitor NU7441 (PKi; 2 μM; n=5; mean±SEM; paired t-test). **(H, I)** As in panels A and B, respectively, but now analyzing total Rad51 intensity per S-phase nucleus (n=4; mean±SEM; one-way ANOVA with post-hoc Dunnett’s). **(J, K)** IF-micropscopy of UV-A laser micro-irradiated cells. Panel J shows representative images, panel K shows the quantification. Plotted are the data from all biological repeats. Each grey or green dot represents an individual track, black dots are the median for each biological repeat (n=4; mean±SEM; ratio paired t-test).

To further interrogate an end-resection function for FANCL, we quantified resected DNA using U-2 OS AsiSI cells. In these cells, the site-directed nuclease AsiSI translocates to the nucleus after tamoxifen treatment and induces DSBs at ∼200 well-defined locations^44,45^. The extent of end-resection can then be quantified as diagrammed in figure 5D^46^. In short, genomic DNA is purified and digested with restriction enzymes that target locations at various distances from a defined AsiSI locus. Unlike dsDNA, resected DNA will remain undigested and can therefore be detected by PCR-amplification using primers flanking the restriction enzyme target site. U-2 OS AsiSI *FANCL*^KO^ and *FANCD2*^KO^ cells were generated and validated by Western blotting for ubiquitinated and total FANCD2 levels (Fig. 5E). The cells were treated with tamoxifen, and end-resection was quantified at sites located 335 bp or 1618 bp downstream from a defined AsiSI locus (Chr 1: 89231183) by qPCR. Resected DNA was detected specifically in the tamoxifen-treated cells, and at a higher frequency at the 335 bp location than the 1618 bp location, consistent with previous reports (Fig. 5F)^46^. Compared to control cells, the frequency of resected DNA was reduced in each of the *FANCL*^KO^ and *FANCD2*^KO^ clones, significantly so at the 1618 bp distance (Fig. 5F). These results are consistent with an end-resection defect in *FANCL*^KO^ and *FANCD2*^KO^ cells, although the phenotype in the AsiSI assay was mild compared to the pRPA phenotype. We therefore repeated the AsiSI assay in the presence of a DNA-PKcs inhibitor (PKi) to block c-NHEJ and direct DSB-repair towards HR. We reasoned that inhibition of canonical end-joining would require maximum end-resection capacity and would therefore more potently reveal a potential end-resection defect. As anticipated, DNA-PKcs inhibition increased the frequency of resected DNA ∼3-fold in control cells, both at the 335 bp location as well as at the 1618 bp location (Fig. 5G). In the *FANCL*^KO^ cells, however, the NU7441-induced increase in end-resection was less pronounced, particularly at the 1618 bp location. Consequently, in the presence of DNA-PKcs inhibitor the frequency of resected DNA was substantially lower in the *FANCL*^KO^ cells compared to the control cells (Fig. 5G). The resection-defect in *FANCL*^KO^ cells in presence of DNA-PKcs inhibitor was also observed when resection was quantified at a second AsiSI locus (Chr 1: 109838221; Fig. S3F). Collectively, these results indicate that FANCL promotes end-resection at DSBs.

To assess the downstream effects of the impaired end-resection in *FANCL*^KO^ cells, we examined the loading of the HR-recombinase Rad51, which is dependent on ssDNA generation. U-2 OS *FANCL*^KO^ cells and their reconstituted controls were exposed to IR, followed by quantification of nuclear Rad51 foci intensity by immunofluorescence microscopy (Fig. 5H, I, S3G). Compared to control cells, the total Rad51 foci intensity per nucleus was significantly reduced in the *FANCL*^KO^ cells expressing either the empty vector or the FANCL LD mutant, but not in the *FANCL*^KO^ cells expressing FANCL WT. Finally, Rad51 accumulation at UV-A laser-induced DNA damage sites was examined, and found to be significantly lower in the *FANCL*^KO^ cells than in wild-type control cells (Fig. 5J, K). Taken together, these data indicate that FANCL promotes Rad51 accumulation at DSBs, consistent with a role for FANCL in end-resection.

### FANCL and Ube2T promote CtIP recruitment to DSBs

Exactly how the FA core complex promotes end-resection at DSBs is unclear. It has previously been shown that FANCD2 directly interacts with the end-resection factor CtIP and is required for its recruitment to MMC-induced interstrand crosslinks^47,48^. We therefore hypothesized that FANCL and Ube2T would similarly promote CtIP recruitment to DSBs. To study this, CtIP accumulation into IR-induced DNA damage foci was quantified in *FANCL*^KO^ and wild-type control cells. Whereas clear CtIP foci were detected in control cells and in FANCL WT reconstituted *FANCL*^KO^ cells, a striking absence of CtIP foci was observed in the *FANCL*^KO^ cells expressing either the empty vector or the FANCL LD mutant (Fig. 6A, B). To further substantiate these findings, CtIP accumulation at UV-A laser-induced DSBs was monitored. CtIP enrichment was reduced at these DSBs in *FANCL*^KO^ cells compared to their wild-type counterparts, and this effect was even more pronounced upon inhibition of DNA-PKcs (Fig. 6C, D). Notably, some CtIP accumulation was still observed at UV-A induced DSBs in the *FANCL*^KO^ cells, indicating that loss of FANCL reduced, but did not completely abrogate CtIP recruitment to DSBs. The complete absence of CtIP foci seen following IR exposure is therefore most likely explained by a reduction in CtIP accumulation in these foci below the fluorescence imaging threshold. Of note, FANCD2-deficiency has been associated with reduced CtIP expression^42^. However, we found no difference in CtIP protein levels between control and *FANCL*^KO^ cells, thus excluding the possibility that FANCL regulates CtIP recruitment indirectly by affecting its expression (Fig. S3H). Instead, our results indicate that FANCL ligase activity is required for optimal CtIP recruitment to DSBs.

**Figure 6.**
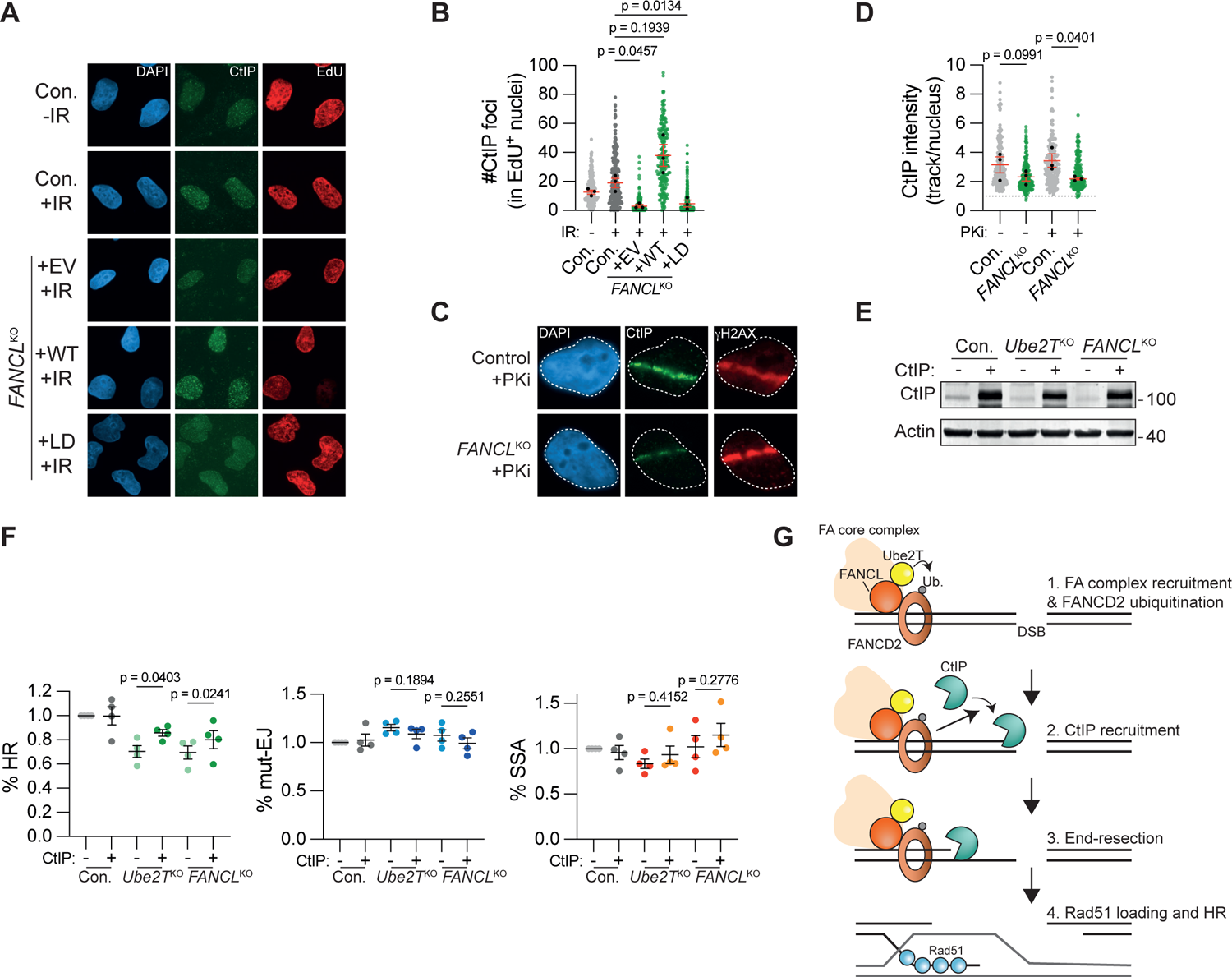
FANCL promotes CtIP recruitment to DNA double-strand breaks. **(A, B)** U-2 OS *FANCL*^KO^ and wild-type control cells were exposed to 10 Gy ionizing radiation (IR), followed by IF microscopy to detect CtIP foci in S-phase (EdU+) nuclei. Panel A shows representative images, panel B shows the quantification. Plotted are the data from all biological repeats. Each grey or green dot represents an individual nucleus, black dots are the median for each biological repeat (n=3; mean±SEM; one-way ANOVA with post-hoc Dunnett’s). **(C, D)** IF-microscopy of UV-A laser micro-irradiated cells. Panel C shows representative images, panel D shows the quantification. Plotted are the data from all biological repeats. Each grey or green dot represents an individual track, black dots are the median for each biological repeat (n=3; mean±SEM; ratio paired t-test). **(E)** Western blot analysis of CtIP overexpression in the cells decribed in panel F. **(F)** Indicated HEK 293T+DSB-Spectrum_V3 cell-lines were transfected to express either CtIP or an empty vector control, together with Cas9 and the sgRNA targeting the BFP gene in the reporter locus. Next, cells were analyzed by flow cytometry to determine the frequency of repair by each of the three indicated pathways. Data were normalized to the Con.+EV (n=4; mean±SEM; One-way ANOVA+Dunnett’s multiple comparison). **(G)** Model depicting how FANCL/Ube2T promotes repair of DSBs by homologous recombination. See main text for details.

Finally, we sought to determine whether the impaired CtIP recruitment was causal to the HR-defect observed in *Ube2T*^KO^ and *FANCL*^KO^ cells. As some CtIP recruitment could still take place in absence of FANCL (Fig. 6C, D), we reasoned that overexpression of CtIP might be sufficient to overcome this recruitment defect. We therefore monitored the frequency of HR repair in control, *Ube2T*^KO^ or *FANCL*^KO^ DSB-Spectrum_V3 cells in the absence or presence of CtIP overexpression (Fig. 6E). While CtIP overexpression did not affect the extent of DSB repair by HR in control cells, it significantly enhanced it in *Ube2T*^KO^ or *FANCL*^KO^ cells (Fig. 6F). In contrast, CtIP overexpression did not significantly affect the levels of DSB repair by mut-EJ or SSA in any of the FA-deficient cell-lines. Hence, CtIP overexpression rescued the HR-defect in FA-deficient cells, indicating that impaired CtIP-dependent end-resection is causal to the reduced HR capacity observed in *Ube2T*^KO^ and *FANCL*^KO^ cells.

## Discussion

Here we report that the FA core complex members FANCL and Ube2T, as well as their downstream substrate FANCD2, promote repair of DSBs through HR. This function is independent of their role in ICL-repair, as FANCL and Ube2T are directly recruited to both UV-A laser- and nuclease-induced DSBs. Their recruitment and activity are required for the accumulation of FANCD2 at DSBs. In the absence of FANCL, Ube2T or FANCD2, DSB end-resection is impaired and Rad51 loading is reduced. This can be explained by reduced recruitment of CtIP, whose accumulation at DSBs is highly dependent on the presence of ligase-proficient FANCL. Collectively, our data support a model in which UbeT2/FANCL drive the ubiquitination and accumulation of FANCD2 at DSBs to promote CtIP recruitment and end-resection during the initial steps of HR (Fig. 6G).

A function for the FA core complex in HR was first suggested approximately two decades ago, based on experiments showing reduced HR in reporter assays in FANCC/FANCG KO chicken DT40 cells^20,49^. These results were later confirmed in mouse and human cells, and extended to other FA-factors, including FANCL and Ube2T^21,22,50–54^. Nevertheless, the importance of the FA core complex and FANCD2 for HR has remained controversial for a number of reasons^55^. Most importantly, the HR-phenotype in reporter assays was generally considered mild, when comparing FA-factor deficient cells to those missing canonical HR factors like BRCA1^21,39,42,55^. Similarly, FA-patient derived cell-lines were only modestly more sensitive to PARPi treatment than wild-type control cells^26,27^. Finally, for FANCA, FANCG and FANCI, the HR-phenotypes were ambiguous^22,28,51^. Here, using an unbiased genetic screening approach, we observed that out of 2,760 genes, FANCL, Ube2T and FANCM were among the most important HR-promoting factors. Similarly, a recently published screen that studied HR with an ectopically provided dsDNA donor, also identified FA genes to be strong drivers of this process^23^. Furthermore, we validated the HR function of FANCL, Ube2T and FANCD2 using orthogonal assays in a variety of isogenic (non-complemented and complemented) knock-out and control cell-line models. These data therefore add compelling evidence for an HR-promoting function of the FA core complex and FANCD2.

We found that HR is not completely abrogated upon depletion of FANCL or Ube2T, despite the use of validated clonal CRISPR KO cells. The residual HR could be explained by FA-factors being required for the repair of a specific subset of DSBs. For example, the FA core complex could be specifically recruited to those DSBs that cause stalling of replication forks, similar to how it is recruited to ICLs upon fork stalling at these lesions^18^. Alternatively, the FA core complex might primarily be required for repair of DSBs with blocked ends. This hypothesis could explain why end-resection of AsiSI-induced DSBs was more dependent on FANCL when DNA-PKcs was inhibited (Fig. 5G). The latter locks the Ku-DNA-PKcs complex on DSB-ends, because auto-phosphorylation of DNA-PKcs is required to loosen its interaction with DNA^56^.

Throughout our studies, we primarily focused on FANCL, Ube2T and FANCD2, but we also showed that FANCD2 accumulation at DSBs is dependent on FANCC. Based on these results, and on published data describing HR-functions for other FA-factors^22,23,57^, we consider it likely that FANCL and Ube2T function as subunits of the complete FA core complex in DSB-repair. During repair of DNA interstrand crosslinks, FANCM plays an important role in recruiting the complex to the lesion^18^. We also identified FANCM as HR-promoting factor in the screen, and it is thus tempting to speculate that it plays a similar lesion-sensing function in DSB-repair. However, *in vitro* FANCM binds most strongly to DNA substrates with branched arms, and interacts poorly with non-branched dsDNA, although some binding to a dsDNA substrate with a 3’ overhang was observed^31^. Since our data indicate that the FA core complex acts upstream of end-resection, it is unlikely that DSBs with 3’ overhangs recruit the FA core complex. Therefore, as an alternative to FANCM, FANCA may function as the lesion-sensing factor, since FANCA was shown to bind to blunt dsDNA in vitro^28^.

Whereas FANCL/Ube2T loss clearly reduced HR, we could not observe any effect on mutagenic end-joining or SSA in our DSB-Spectrum_V3 reporter assays. These end-joining results are in agreement with a study by Howard *et al.*, which reported no effect on total end-joining upon depletion of different FA-factors^22^. However, Howard *et al.*, as well as others, did show a function for the FA core complex and FANCD2 in DSB-repair through a-EJ, in studies using reporters specific for this pathway^22,52,58^. FANCD2 has been shown to promote the accumulation of the a-EJ factor Polε to foci induced by UV irradiation or hydroxyurea^58^. Such an a-EJ function would be consistent with FA-factor dependent recruitment of CtIP, which has been shown to promote a-EJ but not c-NHEJ^59^.

We did not detect an a-EJ phenotype in FA-factor deficient cells in our studies. This can be explained by the observation that the Cas9 target site in DSB-Spectrum_V3 is predominantly repaired through c-NHEJ, and rarely through a-EJ^17^. More surprisingly was the absence of an SSA phenotype in our FANCL/Ube2T deficient cells. Although this observation is in agreement with some other studies^58^, the majority of reporter studies observed a reduction in SSA upon loss of FA-factors^21,22,28^. Impaired SSA would also be expected considering its dependency on end-resection^1^. These discrepant results might be explained by the different reporters used in our studies compared to those used in the other studies. We previously observed that knockdown of all established end-resection factors reduced SSA-repair of DSB-Spectrum_V3, but this effect was relatively mild for CtIP, despite a strong reduction in HR^17^. However, when assessed by the SA-GFP reporter, as used in other studies, CtIP knockdown strongly reduced SSA^22^. For both DSB-Spectrum_V3 and SA-GFP, the SSA-repair product that is measured is a repeat-mediated deletion, but the reporters differ with regards to the length of the repeats (517 bp and 280 bp, respectively) and the distance between repeats (3.2 kb and 2.4 kb, respectively). Furthermore, the DSB is generated by the I-SceI nuclease in SA-GFP, which leaves a four nucleotide 3’ overhang, but by Cas9 in DSB-Spectrum_V3, which generally leaves blunt ends and remains associated with the DNA substrate after nucleolysis^60^. Both these differences between the reporters could affect the requirements put on the end-resection machinery, and hence the extent to which SSA-mediated DSB-repair is dependent on CtIP.

Notwithstanding its function in SSA, CtIP is a core HR-factor that interacts with the MRN-complex^5^, and with BRCA1^10,11^. Perhaps surprisingly, these interactions were demonstrated to be non-essential for CtIP accumulation at DNA lesions^13,14^. In contrast, our data indicate that CtIP accumulation at DSBs is strongly dependent on the FA core complex and FANCD2. Similarly, it has been shown that FANCD2 is required for recruitment of CtIP to ICLs, as well as to DNA damage induced by hydroxyurea or UV irradiation^47,48,58^. Moreover, FANCD2 has been shown to interact directly with CtIP, in a manner dependent on its ubiquitination and on FANCA^47,48^. Thus, a key function of the FA core complex and FANCD2 in genome maintenance appears to be the recruitment of CtIP to sites of DNA damage.

Our findings emphasize that DSB-repair by HR requires the orchestrated activity of a multitude of factors that do not necessarily operate in a linear pathway. These findings furthermore indicate that genome maintenance factors can be involved in multiple pathways, thus causing complex phenotypes when lost or mutated. In line with this notion, inactivating mutations in different multifunctional FA genes cause complex FA disease phenotypes^19^. FA patients, in general, have defective ICL-repair, and an impaired response to replication stress^55^. In addition, our data adds to a body of evidence suggesting that FA-defective cells have a reduced capacity to repair DSBs by HR. Further investigation into the extent to which improper DSB-repair contributes to the FA-disease phenotype is therefore warranted.

## Materials and Methods

### Cloning

Restriction enzymes and T4 DNA ligase, Phusion Polymerase and NEBuilder HiFi DNA Assembly Master Mix were all obtained from New England Biolabs and used according to manufacturer’s instructions. Gel extraction, PCR purification, and DNA mini-, midi- or maxiprepping was done using Qiagen kits according to manufacturer’s instructions. To generate HEK 293T + Cas9 + DSB-Spectrum_V2 cells, a lentiviral Cas9 plasmid was cloned by NheI/BamHI restriction to isolate 3xFLAG-Cas9 from pCW-3xFLAG-Cas9 (Addgene # 50661)^61^ followed by ligation into SpeI/BamHI digested pLVX-IRES-Hygromycin. The pLVX-DSB-Spectrum_V2 plasmid was previously described^17^. To activate the reporter in the genetic screen, pLX-sgRNA (Addgene #50662)^61^ was modified by replacing the blasticidin resistance cassette with mCherry using BspeI/EcoRI based restriction ligation. The BspeI site was blunted with Klenow polymerase (New England Biolabs) prior to ligation. Next, the BFP-targeting sgRNA was cloned into pLX-mCherry using the published protocol (see table 1 for sgRNA sequences)^61^. The sgRNA library was cloned into Lentiguide-Blast, which was generated by replacing the EF1a-Puro cassette in Lentiguide-Puro (Addgene #52963)^62^ with a EF1a-Blasticidin cassette using the Xma/Mlu restriction sites. The CRISPR constructs to make KO cell-lines were generated in pSpCas9-2A-iRFP(670), a previously described derivative of pSpCas9-2A-puro^17^. Cloning of sgRNAs was done by ligating annealed sgRNA primer dimers into BpiI-digested pSpCas9-2A-iRFP(670), as previously described^63^. The sgRNAs were selected from the Brunello sgRNA library^64^ (table 1). To re-express FANCL in the FANCL^KO^ cell-lines, a pLVX-hPGK-FANCL-Hygromycin construct was generated. First the CMV-promoter in the pLVX-Hygromycin lentiviral backbone was replaced with a hPGK promoter by assembling a PCR-derived hPGK fragment into ClaI-digested pLVX-Hygromycin together with FANCL cDNA that was PCR amplified from a cDNA library. For the recruitment assays, GFP-tagged FANCL was cloned by PCR-amplification of eGFP and assembly into XhoI-digested pLVX-hPGK-FANCL-Hygro. GFP-tagged Ube2T was generated in an identical manner, with Ube2T cDNA also derived from a cDNA library. The pSpCas9-BFP-sgRNA-iRFP(670) plasmids for transient DSB-Spectrum_V3 reporter experiments was previously described^17^. The pcDNA-CtIP-2xFLAG construct used for CtIP overexpression was a kind gift from Petr Cejka (Institute for Research in Biomedicine, Bellinzona, Switzerland).

**Table 1.**
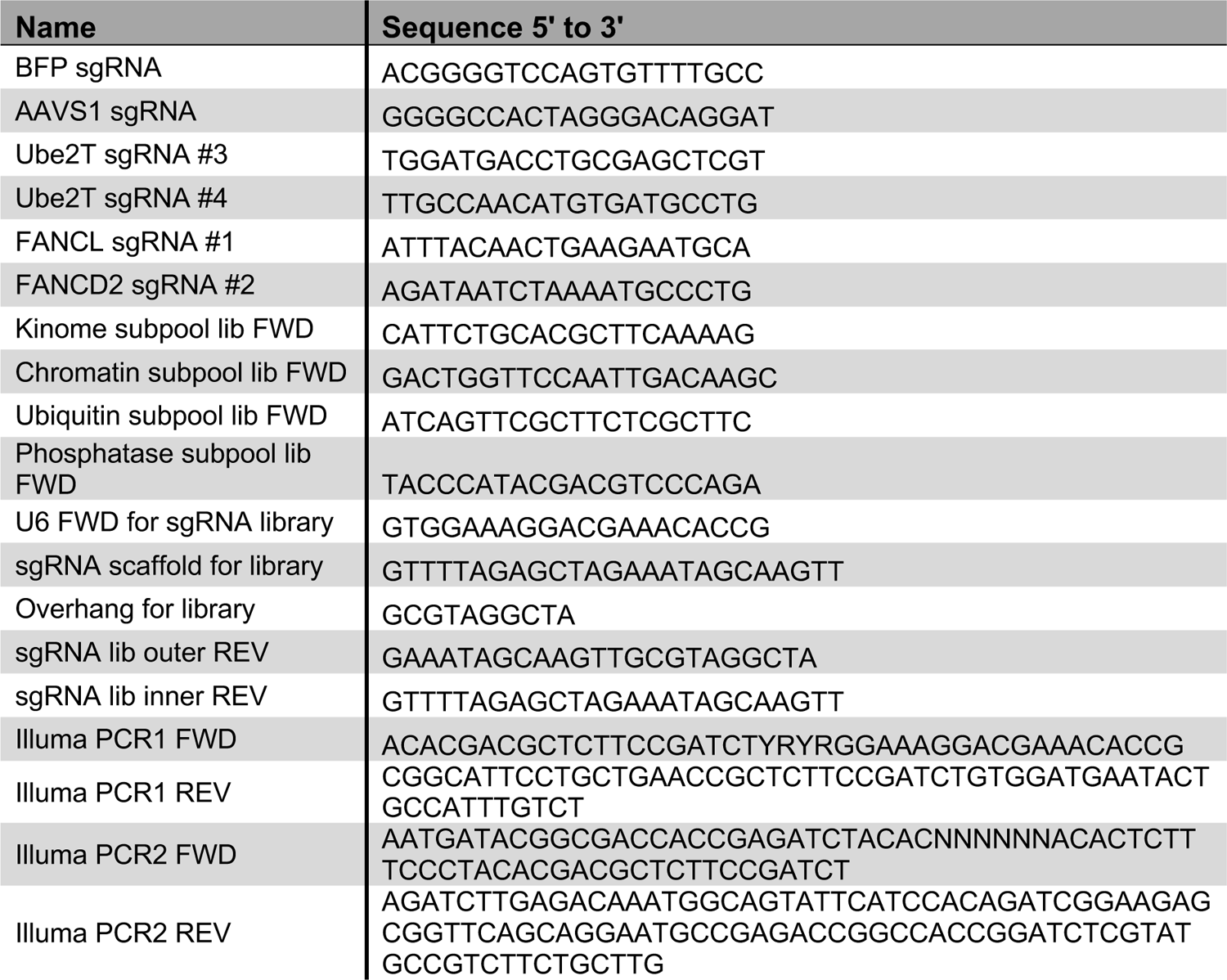
Oligo table.

### Cell-lines

All cell-lines were grown at 37°C with 5% CO_2_ in DMEM supplemented with 10% fetal calf serum and antibiotics. All cell-lines were regularly tested for mycoplasm infection and consistenly found mycoplasm-free. The U-2 OS 2-6-3 cells expressing ER-mCherry-LacI-FokI-DD were a kind gift from Roger Greenberg (University of Pennsylvania, Philadelphia, Pennsylvania, USA)^41^. The U-2 OS AsiSI cells were a kind gift from Gaelle Legube (CBI, Toulouse, France)^45^. The HEK 293T DSB-Spectrum_V3 cell-line was previously described^17^. To generate HEK 293T + Cas9 + DSB-Spectrum_V2 cells, regular HEK 293T cells were lentivirally transduced with pLVX-Cas9-Hygromycin, and selected on Hygromycin B (200 μg/ml; Thermo Fisher Scientific). Next, cells were lentivirally transduced with pLVX-DSB-Spectrum_V2 at low multiplicity-of-infection (MOI) to ensure single integration, and selected on puromycin (1 μg/ml; Invivogen). Subsequently, BFP-positive cells were single-cell sorted by FACS in 96-well plates and expanded. A clone was selected based on the appearance of a sizeable GFP^+^ and BFP^-^ population specifically after reporter activation.

### Lentiviral transduction

To produce lentivirus, HEK 293T cells were transfected with the lentiviral plasmid and packaging plasmids pCMV-VSVg and pCMV-ΔR8.2 at a 6:1:4 mass ratio. Transfection was done using the CalPhos mammalian transfection kit (Clontech). Alternatively, HEK 293T cells were transfected with the lentiviral plasmid and the packaging plasmids pMDLg/RRE, pRSV-REV and pMD2.g at a 4:2:1:1 mass ratio using jetPEI transfection reagent (Polyplus). At 48-72h after transfection, medium with virus was harvested and filtered through a 0.45 μM filter, Next, polybrene (4-8 μg/ml) was added and viral supernatant was added to the recipient cells.

### Generation of the custom sgRNA library

To generate the targeted sgRNA library for the genetic screen, the sgRNA sequences of selected targets were copied from the Brunello library^64^ (see table S1). The targeted library was divided in the following four subpools. Subpool 1 targeted the kinome (763 genes). It already existed as Brunello sub-library and the sequences could be copied directly. Subpool 2 targets chromatin factors (1102 genes). Targets were selected based on association with a chromatin Gene Ontology term, or the presence of a bromo-, chromo-, jumonji, PHD or TUDOR domain, which was determined using AmiGO, Panther and SMART^65–67^. Subpool 3 targeted ubiquitin and SUMO-associated genes (908 genes). This consisted of genes listed in Hutchins *et al*.^68^ supplemented with targets missing from the list but having ubiquitin or SUMO related GO terms, as well as the genes encoding for SUMO or ubiquitin itself. Subpool 4 consisted of phosphatases and targets were selected using the DEPOD database (144 genes)^69^. Furthermore, 350 non-targeting control sgRNAs were added, selected from the Brunello library. The total library therefore contained 12,018 sgRNAs targeting 2760 non-redundant genes, with four sgRNAs per gene.

Oligo’s were designed to contain, from 5’ to 3’, a library specific forward primer sequence (lib FWD), a U6 promoter region for Gibson Assembly (U6 FWD), the sgRNA protospacer sequence, an sgRNA scaffold region for Gibson assembly and a 10 bp overhang (table 1). These 95-96 bp oligo’s were obtained in a pooled format (CustomArray, Inc.), and PCR-amplified using a nested PCR reaction with first the lib FWD primer and an outer reverse primer, followed by a PCR with the U6 FWD and an inner reverse primer. The PCR product was gel-extracted and assembled into BsmBI-digested Lentiguide-Blast using the Quick-Fusion kit (Biotools). Assembled plasmid was transformed by elctroporation into Endura competent cells (Biosearch technologies), followed by DNA prepping of the libraries.

### Genetic screen in DSB-Spectrum_V2 cells

Throughout the experiment the representation of the sgRNA library was maintained at 250x. First, HEK 293T + Cas9 + DSB-Spectrum_V2 cells were lentivirally infected with the sgRNA library at an MOI of 0.2 to ensure single integration. Cells were expanded in the presence of Blasticidin S (10 μg/ml; Invitrogen) to high numbers and thereafter stored in liquid nitrogen.

For each replicate of the screen, sgRNA library infected DSB-Spectrum_V2 cells were taken into culture and allowed to recover from freeze-thawing for several days. Next, they were lentivirally transduced to introduce pLX-BFP-sgRNA-mCherry at a high MOI giving 100% infection and strong sgRNA expression. At seven days after BFP sgRNA infection, 4-12*10^6^ cells of the GFP+, BFP- and total population was harvested by FACS on a BD FACSAria II or III (BD Biosciences). Next, genomic DNA was isolated using the QiaGen Blood and Tissue DNeasy kit according to manufacturer’s protocol. The sgRNA sequences were amplified from the genomic DNA with primers Illumina PCR1 FWD and Ilumina PCR1 REV using Phusion Hot Start Flex polymerase (New England Biolabs), with the annealing temperature set at 58°C. The resulting product was concentrated by PCR purification using the Qiagen PCR purification kit, and subsequently gel-extracted using the Qiagen Gel extraction kit. Next, a short, 12-cycle PCR was performed with Illumina PCR2 FWD and Illumina PCR2 REV to add the Illumina adapters and sequencing primer binding sites. The resulting PCR product was analyzed by agarose gel electrophoresis and gel-extracted. Next, PCR products were sequenced on a HiSeq 2000 sequencing platform (Illumina). A fraction of the sgRNA plasmid library was also sequenced to determine the sgRNA distribution in the input material. Next, sgRNA counts were extracted from the sequencing fastq files. Prior to analysis, any sgRNA with a read-count <50 in the reference population was excluded from all samples in that particular repeat. Finally, sgRNA fold-change and statistical analysis were determined by uploading the read-count files in the BasDAS analysis system^70^, a web-based interface to analyze sgRNA depletion and enrichment using the MAGeCK algorithm^71^. The list of essential genes was obtained from Wang *et al*^30^.

### Generation of knock-out cell-lines

To generate Ube2T/FANCL and FANCD2 knock-out cell-lines, the parental cell-lines were transfected using lipofectamine 2000 (Thermo Fisher Scientific) with pSpCas9-iRFP(670) plasmids containing the relevant sgRNAs. In all cases a control cell-line transfected with an AAVS1-targeting sgRNA was taken along. At 24-72h after transfection, cells were sorted by FACS for high iRFP(670) expression, followed by culturing and expansion for at least seven days. Next, the AAVS1sg control cells were maintained as a pool, and the Ube2T/FANCL and FANCD2 edited cells were single-cell sorted by FACS in 96-wells plates and expanded. Once confluent, growing clones were re-arrayed and split into two 96-wells plates, one for maintenance and one for testing. At 24-72h after replating, the test plate was washed twice with PBS and DirectPCR lysis reagent (Viagen) mixed 1:1 (vol) with ddH_2_O and supplemented with 0.2 μg/ml proteinase K was added to the wells, followed by overnight incubation at 55°C. Subsequently, Proteinase K was inactivated by incubation at 85°C for 1.5h. Next, a PCR using GoTaq G2 polymerase (Promega) was performed to specifically amplify the wild-type target locus, and not any CRISPR-edited loci. Clones for which no PCR product was obtained were considered candidate knock-out cells, and were further expanded. Knock-out status was validated by western blot for Ube2T and FANCD2. For FANCL, the genomic DNA was isolated using the DNAeasy blood and tissue kit (Qiagen), and the target locus was PCR amplified using the FANCL TIDE primers. Next, PCR products were analyzed by Sanger sequencing.

### DSB-Spectrum reporter assays

For the reporter assays using HEK 293T + Cas9 + DSB-Spectrum_V2 cells, the BFP-targeting sgRNA or AAVS1-targeting control sgRNA was introduced by lentiviral transduction with pLX-sgRNA-mCherry. Next, cells were analyzed by flow cytometry on a BD LSRFortessa (BD Biosciences). Gating was done on live cell, single cell, mCherry-positive events, and the frequency of GFP^+^ and BFP^-^ cells was determined. The background GFP^+^ and BFP^-^ frequencies in the AAVS1sg cells were subtracted from those in the BFPsg cells.

For all reporter assays in HEK 293T + DBSDSB-Spectrum_V3 cells, the control cell-line (Con.) was the AAVS1sg-transfected and iRFP(670)-sorted pool (see above). The exception was the experiment to test Ube2TKO clones 3.3 and 4.1. In three of the seven repeats the untransfected parental was used as a control. DSB-Spectrum_V3 cells were transiently transfected with pSpCas-iRFP(670) containing a BFPsg or AAVS1sg. At 48-96h after transfection, cell were harvested and analyzed by flow cytometry, gating was done on live cell, single cell, iRFP(670)-positive events. The frequency of GFP^+^ (HR), BFP^-^/mCherry^+^ (mut-EJ) and BFP^-^/mCherry^-^ (SSA) cells was determined, and corrected for background levels in the respective AAVS1sg population. Next, the frequency of each repair population was divided by the sum of frequencies of all three repair populations. In case of the CtIP overexpression experiment, the resulting values were normalized to the Control+EV population.

### Clonogenic survival assays

Cells were trypsinized and seeded at low densitiy in medium with the indicated concentrations of Olaparib (Selleckchem, S1060). Medium was replaced with fresh medium containing Olaparib 7 days later. At 14 days after plating, cells were washed and colonies were stained with methylene blue (2.5 gr/L in 5% ethanol). Colony number was determined manually, and normalized to the number of colonies in the untreated control plate. Curve-fitting was performed using Graphpad Prism to determine the IC50.

### Western blotting

Cell pellets were lysed on ice in RIPA buffer (50 mM Tris-HCl pH 8.0, 1 mM EDTA, 1% Triton-X100, 0.5% Sodium Deoxycholate, 0.1% SDS, 150 mM NaCl) supplemented with cOmplete EDTA-free Protease Inhibitor Cocktail tablets (Roche), 2 mM MgCl_2_ and Benzonase Nuclease (100 U/ml; Merck Millipore). Insoluble material was pelleted by centrifugation (14,000 rpm, 15 min.), and protein concentration was determined using a BCA assay (Pierce). Next, Laemmli SDS-sample buffer with reducing agent was added to the lysates. Alternatively, cells were harvested and lysed directly in 2x Laemmli sample buffer. Sample were boiled at 95°C for 5 min. Next, protein was separated by SDS-PAGE on a 4-15% Criterion TGX pre-cast midi protein gel (Bio-Rad), and transferred to nitrocellulose membrane using a standard tank electrotransfer protocol. Membranes were blocked with 5% Blotto non-fat dry milk (Santa-Cruz) (W/V) in Tris-Buffered Saline (TBS). Next, primary, as well as secondary antibody staining was performed in Blocking buffer for fluorescent WB (Rockland), diluted 1:1 in TBS with 0.1% Tween-20 (TBS-T). Membranes were imaged on a Odyssey CLx scanner (LI-COR BioSciences), and image analysis was done using ImageStudio (LI-COR BioSciences). All antibodies are listed in table 2.

**Table 2.** Antibody table.

| Target | Species | Company | Cat. No. | Application and dilution |
| --- | --- | --- | --- | --- |
| Ube2T | Rabbit | Abcam | ab140611 | WB (1:1000) |
| FANCD2 | Mouse | Santa Cruz | sc-20022 | IF (1:100) and WB (1:1000) |
| Tubulin | Mouse | Sigma Aldrich | T6199, clone DM1A | WB (1:5000) |
| gH2AX | Mouse | Millipore | #05-636, clone JBW301 | IF (1:2000) |
| MDC1 | Rabbit | Abcam | ab11171-50 | WB (1:1000) |
| pRPA (S4/S8) | Rabbit | Bethyl | A300-245A | IF (1:1000) |
| Rad51 | Mouse | Genetex | GTX70230, Clone 14B4 | IF (1:500) |
| CtIP | Mouse | Sigma Aldrich | MABE1060 | WB (1:1000) and IF (1:1000) |
| CtIP | Mouse | Santa Cruz | sc-271339 | WB (1:500) |
| Actin | Mouse | MpBiomedicals | 869100 Clone C4 | WB (1:1000) |
| BrdU | Mouse | Roche | 11170376001 | FC (1:200) |

### Immuno-fluorescence microscopy

Cells were plated on coverslips and treated 24-48h later. Next, cells were washed with PBS, or with ice-cold CSK buffer (10 mM HEPES (pH 7.4), 300 mM Sucrose, 100 mM NaCl, 3 mM MgCl_2_) in case of Rad51 IF, and pre-extacted with 0.25% Triton-X100 in PBS (or CSK-buffer) on ice for 2 min. Next, cells were fixed with 2% formaldehyde or paraformaldehyde in PBS for 20 min. at room temperature (RT), followed by washing with PBS. Cells were permeabilized again with 0.5% NP-40 in PBS for 5 min. at RT, washed with PBS, and blocked 1% BSA (w/v) in case of Rad51 IF, for 1h at RT. Next, primary and secondary antibody staining was done in PBS + 1% BSA at RT. In all cases, 0.1 μg/mL 4′, 6-diamidino-2-phenylindole dihydrochloride (DAPI) was added to the secondary antibody mixture. If needed, incorporated EdU was labeled by a copper-catalyzed click-it reaction. First, cells were fixed again with 2% formaldehyde for 15 min. at RT. Next, the click-it reaction mix (PBS with 100 mM Trs-HCl (pH 8.5), 1 mM CuSO_4_, 100 mM freshly prepared ascorbic acid) with 1 μl Alexa Fluor™ 647 Azide (Thermo Fisher Scientific) per 5 ml was added to the cells, followed by incubation for 30 min. at RT. Cells were washed with PBS and coverslips were mounted on glass microscopy slides with polymount. Images were acquired on a Zeiss AxioImager M2 wide-field fluorescence microscope with 63x PLAN APO (1.4 NA) oil-immersion objectives (Zeiss).

### UV-A laser micro-irradiation and FokI nuclease assays

For UV-A laser micro-irradiation, cells were plated on coverslips and first treated with 15 μM 5′-bromo-2-deoxyuridine (BrdU) for 24 h to sensitize for the generation of DSBs. Next, growth medium was replaced with Leibovitz’s L15 medium supplemented with 10% fetal calf serum (FCS) and cells were placed in a Chamlide TC-A live-cell imaging chamber. The chamber was mounted on the stage of a Leica DM IRBE widefield microscope stand (Leica) integrated with a pulsed nitrogen laser (16 Hz, 364 nm; Micropoint Ablation Laser System; Andor) that was directly coupled to the epifluorescence path of the microscope and focused through a Leica 40× HCX PLAN APO 1.25–0.75 oil-immersion objective. The laser output power was set to 72–80 to generate strictly localized sub-nuclear DNA damage. Cells were micro-irradiated (two iterations per pixel) using Andor IQ software (Andor). One field of view was micro-irradiated per min., for a time period of fifteen minutes. Directly here-after, cells were pre-extracted and fixed as decribed above. For FokI nuclease assays, U-2 OS 2-6-3 cells stably expressing ER-mCherry-LacI-FokI-DD were plated on coverslips, and treated with 1 μM 4-hydroxy-tamoxifen (4-OHT; Sigma-Aldrich) and 1 μM Shield1 (Clontech lab, now Fisher Scientific) for 4 h. Next, cells were pre-extracted and fixed as described above.

All image analysis was performed using FIJI software (ImageJ). First, the damaged area was identified in the damage marker channel (γH2AX or MDC1) by thresholding to distinguish the laser stripe or FokI focus, and selection using the wand. Next, the signal intensity of GFP-FANCL/GFP-Ube2T or endogenous FANCD2 in the damaged area was determined (I_damage_), as well their average signal in the nucleoplasm (I_nucleoplasm_) and the background signal in the imaged area (I_background_). Subsequently, enrichment was calculated as follows: ((I_damage_ − I_background_)/(I_nucleoplasm_ − I_background_)).

### IR-induced foci analysis

Cells on coverslips were treated with ionizing radiation using a Xylon X-ray generator machine (Y.TU225-D02; 200 KV; 12 mA; dose rate 2 Gy/min), and pre-extracted and fixed 4 h later. The pRPA and CtIP foci were quantified using the Image-J macro “Foci-analyzer” (freely available at https://github.com/BioImaging-NKI/Foci-analyzer; created by Bram van den Broek, the Netherlands Cancer Institute, the Netherlands). The Rad51 foci were analyzed with the Olympus ScanR Image Analysis Software (3.3.0). A dynamic background correction was applied, and single cell nuclei were segmented using an integrated intensity-based object detection module based on the DAPI signal. Rad51 foci segmentation was performed using an integrated spot-detection module to obtain foci counts and foci intensities. Cell cycle staging was performed based on the total DAPI intensity per cell, which scales with DNA content, and the mean EdU-intensity per cell, which indicates DNA synthesis. All downstream analyses were performed on properly detected interphase nuclei containing a 2N-4N DNA content as measured by total and mean DAPI intensities. Fluorescence signal intensities are depicted as arbitrary units.

### AsiSI end-resection assay

The AsiSI end-resection assay was done essentially as described in Zhou *et al*^46^. In short, U-2 OS AsiSI cells were treated with 1 μM 4-OHT, with or without 2μM NU7441 (SelleckChem), and harvested by trypsinization 4 h later. Next, genomic DNA was isolated using the DNeasy blood and tissue kit (Qiagen), and 250 ng DNA was digested with BsrGI (Site 1), BamHI (Site 2) or HindIII (Control; New England Biolabs) overnight at 37°C. Next, a qPCR was performed using 40 ng digested genomic DNA as template with the GoTaq qPCR master mix (Promega) according to manufacturer’s instruction. The qPCR-reaction was run and quantified on a CFX384 C1000 Touch thermal cycler (BioRad). Data were analyzed to obtain the ΔCt for each reaction using BioRad CFX Manager Software version 3.1, and subsequently the % of ssDNA% at each location was calculated as follows: ssDNA% = 1/(2^(△Ct-1) + 0.5)*100.

### Cell-cycle distribution

For cell cycle analysis, cells were pulse labeled with 15 μM BrdU for 1 h, fixed in 70% ethanol, and subsequently denatured in 2 M HCl. Next, cells were stained with α-BrdU antibody and incubated with 0.1 mg/ml propidium iodide, followed by flow cytometric analysis.

### Statistics

All statistical testing was done using Graphpad Prism version 9.5.1 (Dotmatics).

## Acknowledgements

We thank the LUMC Flow cytometry core facility (LUMC, Leiden, the Netherlands) and the Robert A. Swanson (1969) Biotechnology Center (Koch Institute, MIT, Cambridge MA, USA) for assistance. This research was supported by ERC Consolidator (ERC-CoG-617485; H.v.A.) and NWO-VICI grants (VI.C.182.052; H.v.A.), by a fellowship from the Dutch Cancer Society (BUIT 2015-7546; B.v.d.K.), by funding from the Leiden University Medical Center regulation for MSCA-IF Seal of Excellence awardees (B.v.d.K.), by National Institute of Health grants R01-ES015339 (M.B.Y.), R35-ES028374 (M.B.Y.), R01-CA226898 (M.B.Y.), by the joint Cancer Research UK and Brain Tumour Charity funded Brain Tumour Award C42454/A28596 (M.B.Y.), by the Charles and Marjorie Holloway Foundation (M.B.Y.), by the MIT Center for Precision Cancer Medicine, by the Cancer Center Support Grant P30-CA14051 from the National Cancer Institute, by the Center for Environmental Health Sciences Support Grant P30-ES002109 from the National Institute of Environmental Health Sciences (M.B.Y.) and by the Swiss National Science Foundation (310030_197003; M.A.).

---

**Supplementary Figure 1.**
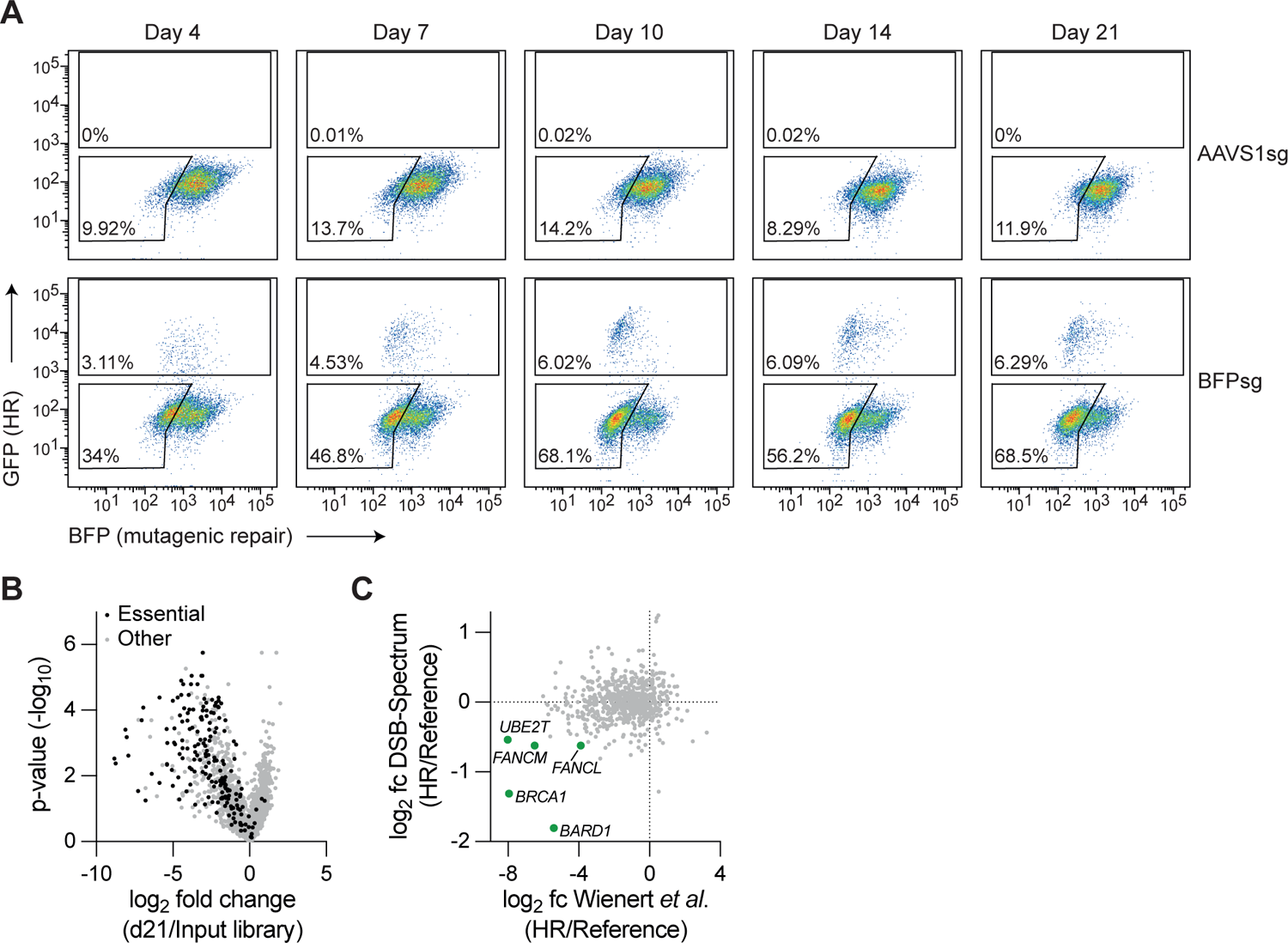
Quality control of a targetd CRISPR screen in DSB-SPectrum reporter cells. **(A)** Representative flow cytometry dot plots of the experiment shown in figure 1B. **(B)** Volcano plot showing the gene targets of sgRNAs that were either enriched or depleted from the total population, as compared to the input sgRNA library, at 14 days after lentiviral introduction of the sgRNA library. Genes were identified as essential based on Wang *et al*., 2015. **(C)** Comparison between this study and Wienert *et al*., 2020 of sgRNA depletion and enrichment in the HR population compared to the reference population. HR-promoting factors that were identified by both studies are indicated in green (fc=fold change).

**Supplementary Figure 2.**
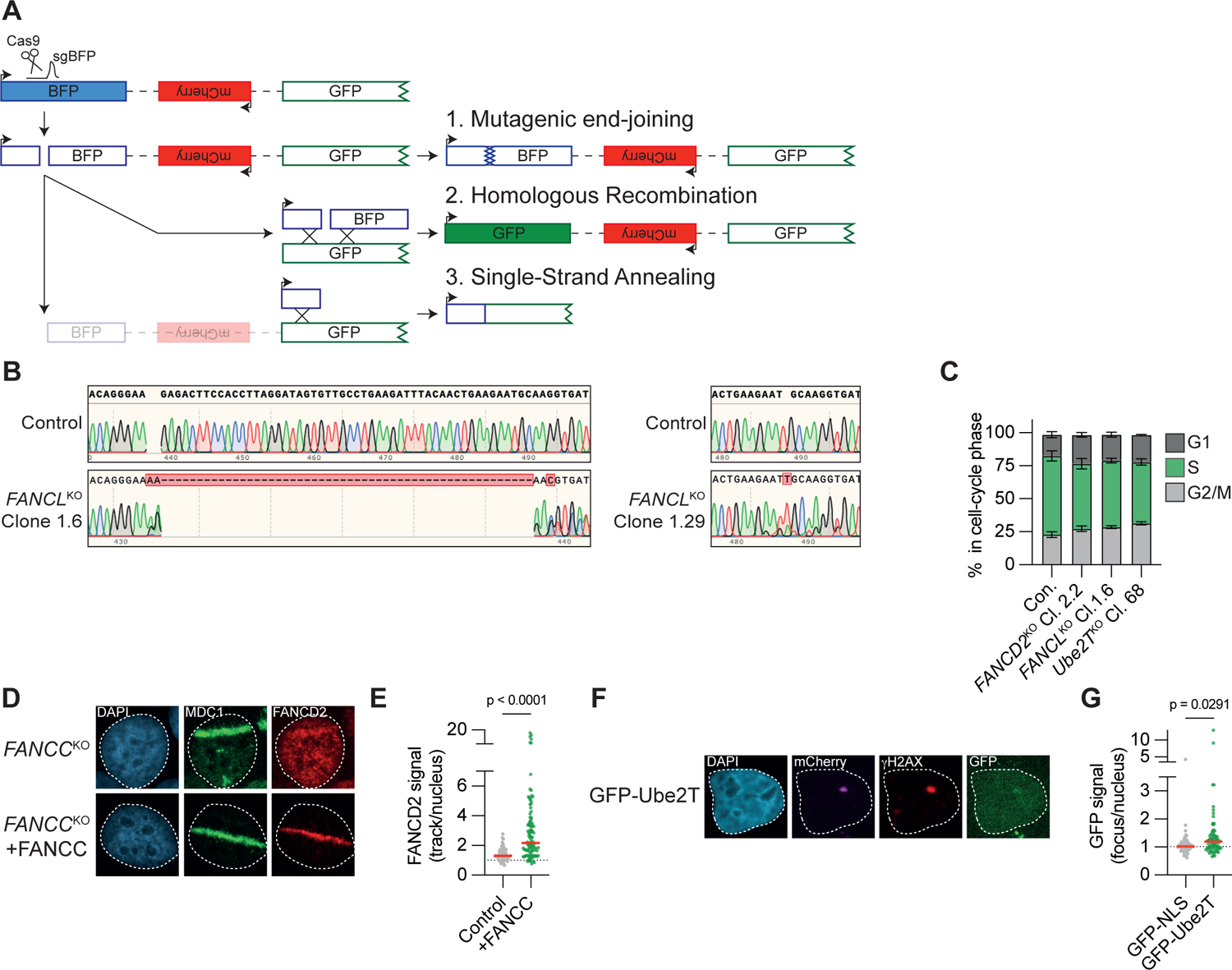
Supporting data to main figures 2, 3 and 4. **(A)** Schematic of the DSB-Spectrum_V3 reporter. Adapted from van de Kooij *et al*., 2022. **(B)** DNA sequence alignment of the *FANCL* sgRNA target site in unedited control cells and the U-2 OS *FANCL*^KO^ clones. Depicted are representative sequence chromatograms, red shaded boxes indicate deviations in the DNA sequence of the FANCLΔ clone compared to control. **(C)** The cell-cycle profile of indicated, asynchronously growing, cell-lines was determined by flow cytometry. The differences between each of the knock-out cell-lines and the wild-type control cell-line were not significant, for neither of the cell-cycle stages (n=3; mean±SEM; One-way ANOVA+-Dunnett’s multiple comparison; p-value ranged from 0.1009 to 0.5659). **(D, E)** FANCD2 recruitment to UV-A laser induced DNA damage was analyzed by imunofluorescence microscopy in FANCC-deficient VU1131 fibroblasts, reconstituted or not with FANCC cDNA. Panel C shows representative images, panel D shows the quantification of a representative experiment. Dotted line is set at 1 (*i.e.* no recruitment to the track), red lines indicate median (n=2; Mann-Whitney test). **(F, G)** As in figure 4F, but for GFP-Ube2T. Panel E shows representative images, panel F shows the quantification of a representative experiment. Red lines indicate the median (n=2; Mann-Whitney test).

**Supplementary Figure 3.**
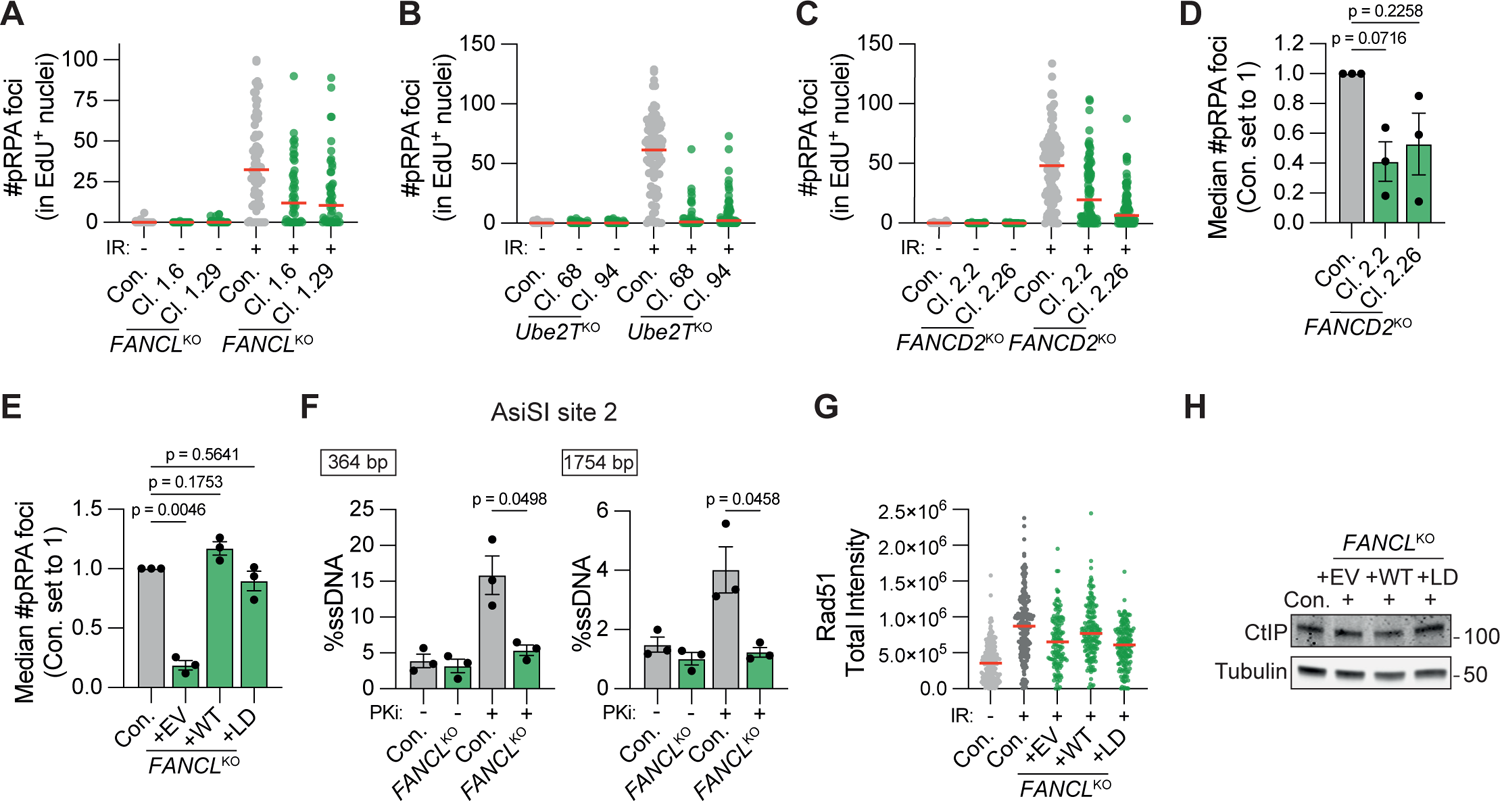
FANCL, Ube2T and FANCD2 promote end-resection at DNA double-strand breaks. **(A)** Depicted is a representative biological repeat of the experiment shown in figure 5A, B. **(B)** As in figure 5A,B, but for *Ube2T*^KO^ cells. Depicted is a representative experiment of two biological repeats. **(C, D)** As in figure 5A/B, but for *FANCD2*^KO^ cells. Panel A shows the quantification of a representative experiment, panel D shows the quantificaiton of the three bioligical repeats (n=3; mean±SEM; one-way ANOVA with post-hoc Dunnett’s). **(E)** For the experiment shown in figure 5C, the number of pRPA foci per S-phase nucleus was plotted (n=3; mean±SEM; one-way ANOVA with post-hoc Dunnett’s). **(F)** As in figure 5F, but for a second AsiSI-targeted locus (n=3; mean±SEM; paired t-test). **(G)** Depicted is a representative biological repeat of the experiment shown in figure 5H. **(H)** Western blot showing CtIP protein levels in control cells and reconsituted U-2 OS *FANCL*^KO^ cells lines.

**Supplementary Figure 4.**
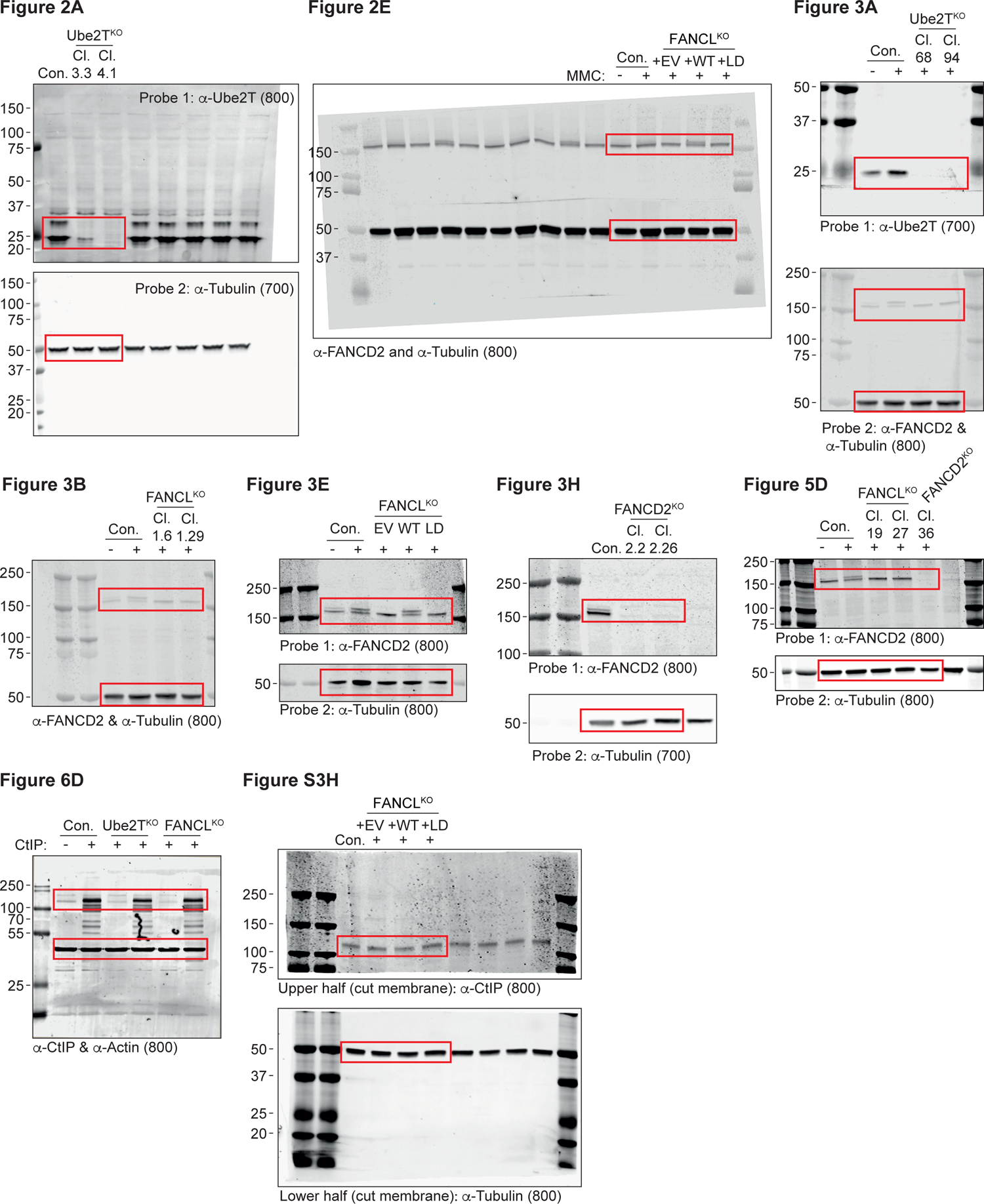
Uncropped western blots. Depicted are all completely uncropped western blot images. Note that in most cases nonly the relevant region of the western blot membrane was imaged, rather than the whole membrane. Red boxes indicate the area selected for the cropped images. The (700) and (800) indications refer to the Oddyssey channel used for imaging.

